# A test of the gleaner-opportunist trade-off among photosynthetic traits in Cryptophyte algae

**DOI:** 10.1101/2023.03.15.530972

**Authors:** Jake A. Swanson, Matthew J. Greenwold, Tammi L. Richardson, Jeffry L. Dudycha

## Abstract

As photosynthetic producers, phytoplankton form the foundation of aquatic food webs. Understanding the relationships among photosynthetic traits in phytoplankton is essential to revealing how diversification of these traits allow phytoplankton to harvest energy from different light environments. We investigated whether the diversification of 15 species of cryptophytes, a phylum of phytoplankton with diverse light-capturing pigments, showed evidence of trade-offs among photosynthetic performance traits as predicted by the gleaner-opportunist resource exploitation framework. We constructed photosynthesis vs. irradiance (P-E) curves and rapid light curves (RLCs) to estimate parameters characterizing photosynthetic performance and electron transport rate. We inferred the evolutionary relationships among the 15 species with ultraconserved genomic elements and used a phylogenetically controlled approach to test for trade-offs. Contrary to our prediction, we observed a positive correlation between maximum photosynthetic rate, *P_max_*, and *P*-*E α*, an indicator of a species’ sensitivity to increases in light intensity when light is a scarce resource. This result could not be explained by electron transfer traits, which were uncorrelated with photosynthetic rate. Together, our results suggest that ecological diversification of light exploitation in cryptophytes has escaped the constraints of a gleaner-opportunist tradeoff. Photosynthetic trade-offs may be context or scale dependent, thereby only emerging when investigated in situations different from the one used here.

## Introduction

Trade-offs among traits can limit evolutionary diversification while simultaneously promoting ecological diversity (Buckling et al., 2003; Kneitel & Chase 2004; Blanchard & Moreau, 2016). Trade-offs constrain diversification by limiting the combinations of traits that are available to an organism, thereby restricting its ability to adapt to a novel environment. Conversely, trade-offs promote ecological diversity by preventing the emergence of a “Darwinian demon,” i.e., an organism that is optimally adapted to its environment in all respects and therefore outcompetes all other species in its community (Kneitel & Chase, 2004). Resource allocation trade-offs have been extensively studied because they are expected to strongly influence patterns of phenotypic variation. There has been less attention to trade-offs in resource acquisition and assimilation, however, even though these processes provide the foundation for allocation decisions and can confound predictions about allocation trade-offs (Van Noordwijk & de Jong, 1986). What work has been done has largely addressed consumers (Llandres et al., 2011), which potentially overlooks the importance of resource acquisition and assimilation trade-offs for primary producers.

The gleaner-opportunist trade-off describes a trade-off between maximum growth rate and minimum resource requirement, R* (Grover, 1990; Bernhardt et al., 2020). A gleaner grows relatively well at low resource levels but relatively poorly at high resource levels (Figure 1a). In contrast, an opportunist grows relatively poorly at low resource levels but relatively well at high resource levels (Figure 1a). A gleaner, therefore, has a lower R* value for the resource while an opportunist has a higher per-capita maximum growth rate. A lower R* value indicates the ability for a population to persist at low levels of a resource, whereas a higher per-capita maximum growth rate indicates the ability to dominate a community when resources are abundant. Later, the gleaner-opportunist framework was extended to include species-specific mortality rates (Litchman & Klauismeier, 2001).

**Figure 1:**
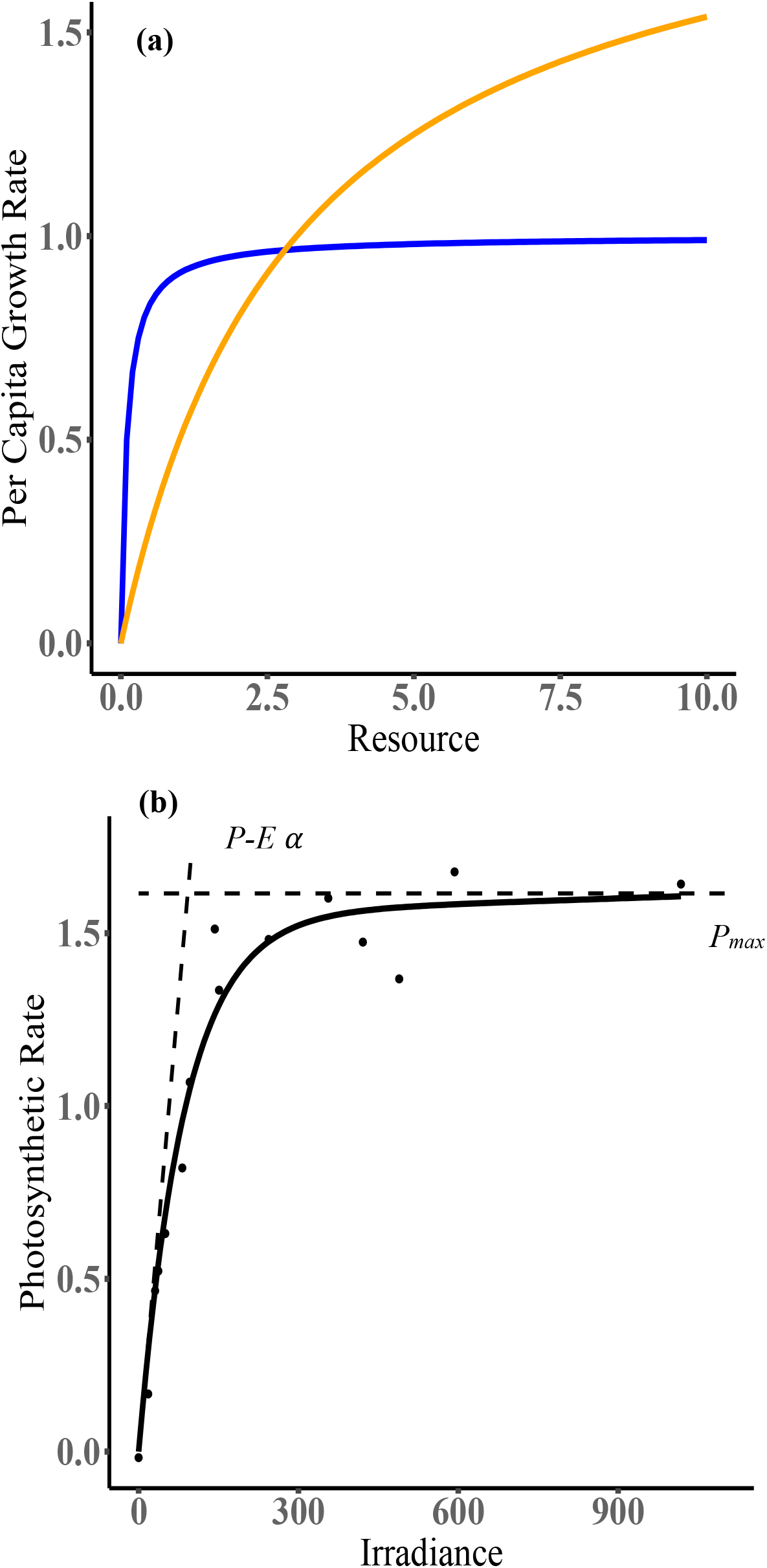
a) Visualization of the gleaner-opportunist framework, adapted from Litchman & Klausmeier, 2001. A gleaner (blue line) has a higher per-capita growth rate at low resource concentrations while an opportunist (gold line) has a higher per-capita growth when resources are abundant. b) Example photosynthesis vs. irradiance curve for the cryptophyte *Rhodomonas salina* indicating the initial slope of the P-E curve, *P-E α*, and the maximum photosynthetic rate, *P_max_*

Gleaner-opportunist trade-offs are believed to be important in the maintenance of diversity in ecological communities (Litchman et al., 2007; Yamamichi & Letten, 2022). The underlying assumption is that this trade-off allows for the coexistence of multiple species when they are competing for a variable resource. Recent work, however, has called into question the existence and relative importance of the gleaner-opportunist trade-off in structuring communities (Kiørboe & Thomas, 2020; but see Letten & Yamamichi, 2021).

Here, we envision phytoplankton as consumers of light, and apply the gleaner-opportunist framework to the photosynthetic and physiological dynamics of light capture and exploitation. We take the view that evolutionary diversification is an important structuring force in ecological communities (McPeek, 1996). Thus, our aim is to test the predictions of this resource acquisition and assimilation framework as applied to the diversification of photosynthetic traits.

### Phytoplankton and photosynthesis

Phytoplankton form the foundation of most aquatic food webs (Field, 1998), thereby determining how much energy is available and influencing diversity at higher trophic levels. Phytoplankton show substantial variation of light capture and photosynthetic abilities. This includes variability of photosynthetic rates (Glover et al., 1987), of the wavelengths of light they can capture (Stomp et al., 2004), and of competitive abilities for light when light intensity fluctuates (Guislain et al., 2019). Light capture and subsequent photosynthetic performance can be considered key resource acquisition and assimilation traits (Richardson et al., 1983).

To investigate a potential gleaner-opportunist trade-off between photosynthetic traits, we performed a phylogenetically controlled analysis across a phylum of phytoplankton with diverse light capture abilities, the Cryptophyta. Cryptophytes are ideally suited to testing general predictions about photosynthetic trade-offs because they encompass substantial photosynthetic diversity, they are monophyletic, and they are commonly found in environments where competition for light may be strong. Cryptophytes’ diverse light capture abilities stem from evolutionary diversification of their pigmentation (Doust et al., 2006; Greenwold et al., 2019). Cryptophyte light capture pigments include chlorophyll *a*, chlorophyll *c_2_*, and eight phycobiliproteins that are unique to cryptophytes (Hoef-Emden & Archibald, 2016). As a taxon, cryptophytes are defined as a monophyletic group by a unique secondary endosymbiosis event (Hoef-Emden & Archibald, 2016). Most other photosynthetic taxa have limited pigment diversity (if any at all), and analyses at higher taxonomic scales are complicated by reticulate evolution. Taken together, these characteristics of cryptophytes allow for comparative investigation of photosynthetic traits while controlling for shared evolutionary history. Furthermore, as frequent inhabitants of low-light environments, cryptophytes face strong pressure to optimize photosynthetic capabilities, making them a good model for investigating adaptive hypotheses about photosynthetic diversification. Other photosynthetic organisms that live in low-light environments such as shade-tolerant trees in northern hardwood forests (Walters & Reich, 1996), understory plants in forests (Craine & Dybzinski, 2013; Onoda et al., 2015), and coral symbionts (Anthony & Hoegh-Guldberg, 2003) lack either known pigment diversity or monophyly.

We estimated cryptophyte photosynthetic parameters using photosynthesis vs. irradiance curves (P-E curves; “E” is the standard symbol for irradiance in photophysiology; Kirk, 1994) and rapid light curves (RLCs). P-E curves provide an estimate of an organism’s maximum photosynthetic rate, *P_max_*, when light is abundant, along with information about how rapidly rates of photosynthesis rise with increasing light intensity at low levels (Figure 1b). The initial slope of a P-E curve, *α* (here referred to as *P-E α*), is the rate of photosynthesis per unit biomass per unit of incident light (Figure 1b). *P-E α* measures how effectively an organism responds to light at sub-saturating intensities (Kirk, 1994).

RLCs provide information about photosynthesis on a short time scale and provide measurements of relative electron transport rate (*rETR*), and effective quantum yield (Ralph & Gademann, 2005). Effective quantum yield is the proportion of absorbed photons that are used to drive electrons through photosystem II, whereas *rETR* is calculated by multiplying the quantum yield of photosynthesis by the photosynthetically active radiation. RLCs also provide estimates of the maximum relative electron transport rate, *rETR_max_*, between photosystems II and I during the light-dependent reactions, and of the initial slope of a RLC (here referred to as *RLC α*) (Ralph & Gademann, 2005). *RLC α* is an estimate of the sensitivity of electron transfer to variation of light intensity at sub-saturating intensities (Ralph & Gademann, 2005) whereas *rETR_max_* is analogous to *P_max_;* it describes the maximum rate of electron transfer between photosystem II and photosystem I under saturating light. As electron transport underpins photosynthesis more broadly, measuring electron transport traits allows us to investigate mechanisms in more detail than P-E curves.

### Terminology and hypotheses

One challenge for applying the gleaner-opportunist trade-off framework to photophysiology is that the use of photosynthetic traits and terminology may cloud any discussion. For our study, maximum photosynthetic rate and maximum electron transport will take the place of maximum growth rate in the gleaner-opportunist framework. In the photosynthesis literature, prior authors have called *P-E α* and *RLC α* “efficiency” (Kirk, 1994; Ralph & Gademann, 2005), which may cause confusion with other uses in the literature on the ecology of resource exploitation (Watt, 1986; Raubenheimer & Simpson; 1996, Tessier et al., 2000). To avoid this confusion, we will use “sensitivity” to light intensity to refer to the biological interpretation of both *P-E α* and *RLC α*. Additionally, we have physiological rather than demographic data for our species, so we define a gleaner as one that performs better at low light levels but worse at high light levels (greater *P-E α*/*RLC alpha* but lower *P_max_*/*rETR_max_*,) whereas an opportunist performs better at high light levels but worse at low light levels (lower *P-E α*/*RLC α* but higher *P_max_*/*rETR_max_*,).

We hypothesized, that if a gleaner-opportunist trade-off exists, it would manifest as a negative linear relationship between *P_max_* and *P-E α*. Given that we found the opposite (see below), we hypothesized that this could be explained by a trade-off between the two parameters describing electron transport, *rETR_max_* and *RLC α*. Lastly, after examining both P-E curve and RLC data, we tested whether there was a positive relationship between photosynthetic rates and relative electron transport rates, predicting that greater electron transport would be correlated with a greater rate of carbon fixation.

## Methods

### Culture conditions

We used 15 species of cryptophytes in this experiment, 14 of which were obtained from culture repositories (Appendix S1: Section S1). We chose species to represent a wide range of taxonomic, phylogenetic, and functional diversity. Each of the eight unique cryptophyte phycobilins are represented by at least one species in our dataset. We isolated one species from Congaree National Park, Congaree, SC, USA. Of the 15 species, 11 are identified to the species level, two to the genus level, while one, our new field isolate, is an undescribed species of *Cryptomonas* (Greenwold et al., unpublished data) Nine species are marine and six are freshwater. Additionally, we used *Goniomonas avonlea* (CCMP 3327), a non-photosynthetic cryptophyte, as an outgroup for phylogenetic estimation. More detailed culture information can be found in Appendix S1: Section S1.

### Photosynthesis vs. irradiance (P-E) curves

A modified method of Lewis and Smith (1983) was used to measure photosynthesis as a function of irradiance (P-E). In short, NaH^14^CO_3_ was added to a 20mL sample of each species, taken when cultures were growing at mid-exponential phase, to achieve a final activity of approximately 3 μCi mL^-1^. One mL of sample was then dispensed into each of 16 scintillation vials and vials were exposed to light intensities ranging from 0 - 1400 μmol photons m^-2^ s^-1^ and irradiance was measured with a quantum scalar irradiance sensor (Biospherical Instruments, Inc., San Diego, CA, USA) inserted into an empty scintillation vial. Samples were incubated with ^14^C for 20 minutes. After incubation, samples were terminated with 50 μL buffered formalin while dissolved inorganic carbon was driven off by adding 200 μL of 50% HCl and shaking the open vials for at least 12 h (usually overnight) in a fume hood. Five mL of scintillation cocktail (EcoLumeTM; MP Biomedicals, Solon, OH, USA) were then added to each vial, vials were mixed, and radioactive decay was counted using a Beckman LS-6500 scintillation counter (Beckman Coulter Inc., Brea, CA, USA). Photosynthetic rate for each species was measured within a four-hour window (between 14:00 and 18:00 each day). Disintegrations per minute were then converted to chlorophyll *a*-specific primary productivity (Knap et al., 1996). Chlorophyll-specific rates of photosynthesis were plotted against light intensity and curves were fit with the equation of Platt et al. 1980 (Appendix S1: Equation S1). *P_max_* and *P-E α*, were calculated following Platt et al. 1980 (Appendix S1: Equation S2). More details about the P-E curve methods can be found in Appendix S1: Section S1.

### Rapid light curves and electron transport activity

Parameters of electron transport for each species were assessed via the generation of RLCs using pulse-amplitude-modulated (PAM) chlorophyll *a* fluorescence with a Walz Water-PAM (PAM, Heinz-Walz, Germany). 20mL samples of each species, again taken at mid-exponential phase, were dark-adapted for 20 minutes and then 3mL sub-samples of each species were exposed to nine pulses of actinic light increasing in intensity with a range from 30 to 1300 μmol photons m^-2^s^-1^ with a 30 second interval between each light pulse. Curves were fit like Equation S1, with 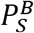 being replaced by *rETR_mPot_* and *P^B^* being replaced by *rETR* (Appendix S1: Equation S3). Estimates of *rETR_max_* and *RLC α* were calculated in the same manner as *P_max_* and *P-E α* (Appendix S1: Equation S4). Electron transport traits were measured in triplicate for each species. More details about the RLC methods can be found in Appendix S1: Section S1.

### Phylogenetic estimation and comparisons across species

Any relationships among photosynthetic traits in cryptophytes may be influenced by the species’ shared evolutionary history, a problem that can be avoided with phylogenetic comparative approaches (Felsenstein, 1985). Therefore, we used phylogenetic generalized least squares (PGLS) to control for phylogenetic history when testing for trade-offs (Martins & Hansen, 1997; Mundry, 2016). We first needed to reconstruct the evolutionary history of our species, which we did with ultraconserved elements (UCEs; Faircloth et al., 2012; 2015). This approach provides a genome-wide perspective on evolutionary relationships with thousands of loci, providing clear benefits over work with only one or two loci (Pamilo & Nei, 1988).

We extracted DNA from each of our species and sent DNA samples to RAPiD Genomics, LLC (Gainesville, Florida) where target enrichment sequencing was performed using Illumina 2 × 150 bp reads. We used RAxML version 8.0.19 for phylogenetic inference (Stamatakis, 2014) with sequence data from 1,868 conserved nuclear genome loci. Details of UCE probe design, DNA extraction and sequencing, and phylogenetic inference can be found in the supplementary material (Appendix S1: Section S1).

### Statistical analysis

Photosynthetic and electron transport trait values were estimated with a non-linear least squares method using the *minpack.lm* package (Elzhov et al., 2016).We performed PGLS analyses in R version 3.6.2 (R Core Team, 2020) using the package *caper* (Orme et al., 2018). We tested for correlations between *P_max_* and *P-E α* and mean *rETR_max_* and mean *RLC α*. An analysis comparing *P_max_* and *rETR_max_* was also done post-hoc after examination of the P-E curve and RLC data. PGLS models were run with habitat, phycobiliprotein absorption peak, and cell volume as predictor variables. These were all non-significant predictors for all models (Appendix S1: Table S3), therefore all models are shown using only the single variable of interest as a predictor (Table 2). We used FigTree version 1.4.0 (https://github.com/rambaut/figtree/releases) to produce the phylogenetic diagram (Figure 2), rooting the phylogeny with *Goniomonas avonlea* (CCMP3327). All other figures were created using *ggplot2* (Wickham, 2016).

**Table 1:** Values for photosynthetic performance traits and electron transport rate traits for each cryptophyte species

| Species | $P-E \alpha$ ($\mu\text{gC } \mu\text{Chl } a^{-1} \text{ h}^{-1}$ ($\mu\text{mol photons m}^{-2} \text{ s}^{-1}$)<sup>-1</sup>) | $P_{max}$ ($\mu\text{gC } \mu\text{Chl } a^{-1} \text{ h}^{-1}$) | $RLC \alpha$ ($\mu\text{mol electrons photons}^{-1}$) | $rETR_{max}$ ($\mu\text{mol electrons m}^{-2} \text{ s}^{-1}$) |
| --- | --- | --- | --- | --- |
| *Cryptomonas ovata* | 0.030 | 3.07 | 0.323 | 58.86 |
| *Chroomonas mesostigmatica* | 0.065 | 2.84 | 0.205 | 21.28 |
| *Chroomonas nordstedtii* | 0.0035 | 0.072 | 0.297 | 35.86 |
| *Chroomonas placodea* | 0.023 | 2.67 | 0.194 | 21.83 |
| *Chroomonas sp.* | 0.015 | 0.24 | 0.222 | 20.78 |
| *Cryptomonas sp.* | 0.048 | 1.93 | - | - |
| *Guillardia theta* | 0.024 | 3.19 | 0.301 | 67.06 |
| *Hemiselmis cryptochromatica* | 0.11 | 2.66 | 0.256 | 27.06 |
| *Hemiselmis pacifica* | 0.031 | 1.79 | 0.356 | 16.96 |
| *Hemiselmis tepida* | 0.030 | 1.43 | 0.212 | 19.04 |
| *Proteomonas sulcata* | 0.058 | 3.05 | 0.367 | 29.31 |
| *Rhodomonas minuta* | 0.061 | 3.47 | 0.323 | 89.66 |
| *Rhodomonas salina* | 0.018 | 1.56 | 0.374 | 30.29 |
| *Teleaulax sp.* | 0.039 | 2.24 | 0.374 | 77.74 |
| *Unid. sp.* | 0.10 | 7.10 | 0.328 | 35.89 |

**Table 2:** Summary of results from PGLS regression models. Significant *p*-values are in bold.

| PGLS Model | Intercept | $F$ | Df | Adjusted $R^2$ | $p$ | N | Pagel's $\lambda$ |
| --- | --- | --- | --- | --- | --- | --- | --- |
| $P_{\max} \sim P\text{-}E_{\alpha}$ | 0.88 | 12.12 | 1,13 | 0.44 | **0.0041** | 15 | 0.054 |
| $rETR_{\max} \sim$<br>$RLC_{\alpha}$ | 20.66 | 0.53 | 1,12 | -0.038 | 0.48 | 14 | 1 |
| $P_{\max} \sim rETR_{\max}$ | 2.64 | 0.038 | 1,12 | -0.080 | 0.85 | 14 | 1 |

**Figure 2:**
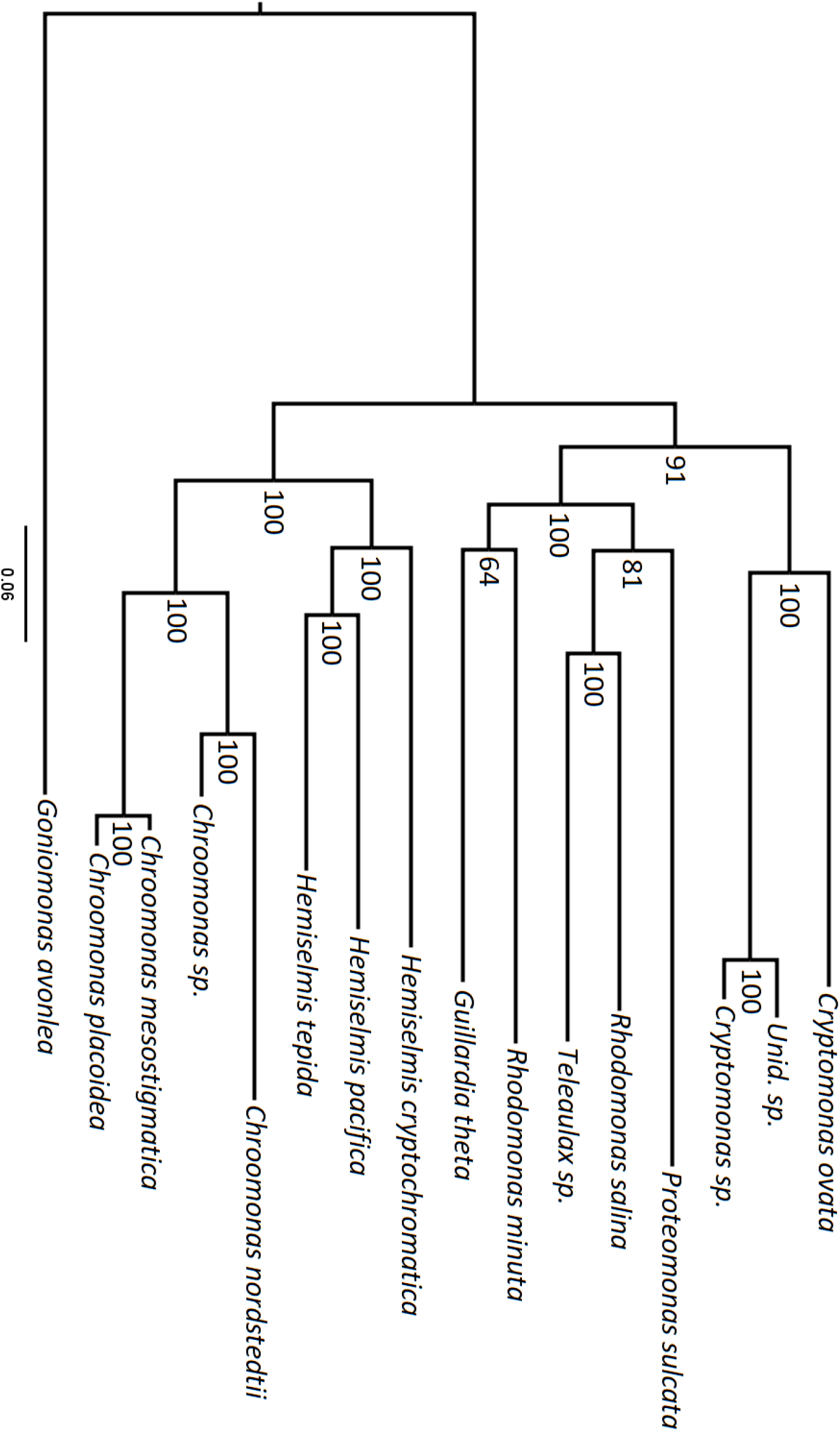
Phylogeny for the 15 cryptophyte species constructed using ultra-conserved elements. Bootstrap values shown at each node.

## Results

### Photosynthetic parameter estimates

Our estimates for *P_max_* spanned two orders of magnitude, ranging from 0.071-7.10 μgC μChl^-1^ h^-1^. Estimates for *P-E α* almost spanned two orders of magnitude, ranging from 0.0035-0.11 μgC μChl^-1^ h^-1^ (μmol photons m^-2^ s^-1^) (Table 1). Variation of photosynthetic parameters estimated from the RLCs were constrained to less than one order of magnitude, ranging from 16.96-89.66 μmol electrons m^-2^s^-1^ for *rETR_max_* and 0.19-0.37 μmol electrons photons^-1^ for *RLC α* (Table 1). We also compared our estimates to photosynthetic parameter values estimated by the Marine Primary Production: Model Parameters from Space (MAPPS) project, which contains estimates of photosynthetic parameters from a global set of 5,711 P-E experiments with marine phytoplankton (Bouman et al., 2018). Our estimates for *P_max_* and *P-E α* for the 15 species of cryptophytes used in this study are in the typical range of estimates from the MAPPS project, indicating that cryptophytes do not have extreme photosynthetic traits (Appendix S1: Section S2, Figure S2).

### Cryptophyte phylogeny

We inferred a phylogeny (Figure 2) with two main clades: 1) a *Hemiselmis*/*Chroomonas* clade, and 2) a *Cryptomonas*/*Rhodomonas* clade. All nodes were strongly supported. To our knowledge, the phylogeny presented here is the first to apply a broad genome-wide data set to Cryptophyta.

### Tests of trade-offs between photosynthetic traits

*P_max_* and *P-E α* are positively correlated (Figure 3a, Table 2) with the correlation having a negligible phylogenetic signal (Table 2). We found no evidence of a correlation between the maximum relative electron transport rate, *rETR_max_* of a species and the initial slope of its RLC, *RLC α* (Figure 3b, Table 2). There was, however, a strong phylogenetic signal (Table 2). We did not find evidence of a correlation between *P_max_* and *rETR_max_* (Figure 3c, Table 2) however a phylogenetic signal was detected (Table 2).

**Figure 3:**
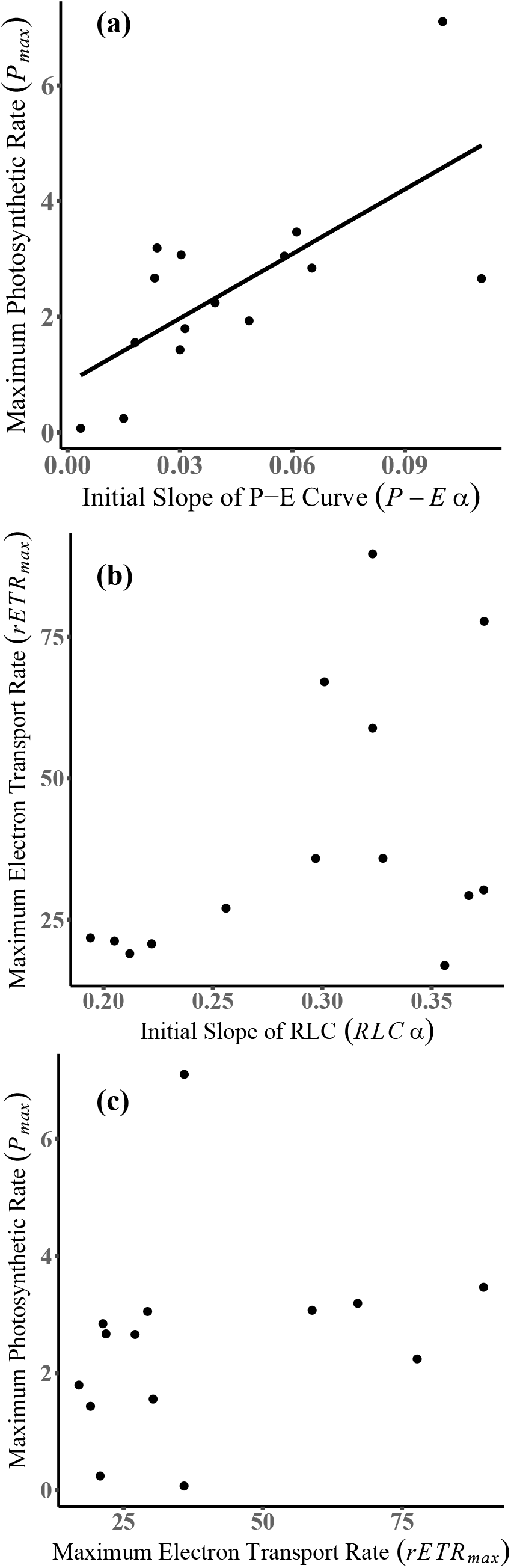
Relationships between photosynthetic parameters derived from P-E curves and RLC. a) Maximum photosynthetic rate, *P_max_* (μg C (μ chl *a*)^-1^ h^-1^), of a species and *P-E α* (μg C (μchl *α*)^-1^ h^-1^ (μmol photons m^-2^ s^-1^)^-1^, the initial slope of its P-E curve. b) Maximum relative electron transport rate, *rETR_max_* (μmol electrons m^-2^s^-1^) and the initial slope of a RLC, *RLC α* (electrons photons^-1^). Points are the mean values of triplicate estimates for *rETR_max_* and *RLC α* for each species or species. c) No correlation between the maximum photosynthetic rate of a species, *P_max_*(μg C (μchl *a*)^-1^ h^-1^), and its maximum relative electron transport rate, *rETR_max_* (μmol electrons m^-2^s^-1^). *rETR_max_* values represent the mean of triplicate estimates for each species or species.

## Discussion

### No evidence of a gleaner-opportunist trade-off

We investigated whether diversification of cryptophyte algae shows a trade-off between photosynthetic traits in the context of a gleaner-opportunist framework. We accounted for evolutionary history in our analyses, allowing us to exclude evolutionary history as a driver of the presence or absence of correlations among traits.

We found no evidence of a gleaner-opportunist trade-off between photosynthetic performance traits in cryptophytes. In fact, we found the opposite: a significant positive correlation between *P_max_* and *P-E α* (Figure 3a). Cryptophytes that respond better to variation of light at low levels also perform better at high light levels, suggesting that photosynthesis is optimized simultaneously across a broad range of light intensities. All species were exposed to light intensities that ranged from 0-1400 μmol photons m^-2^ s^-1^, which reflect light intensity levels that marine and freshwater algae encounter in natural environments (Kirk, 1994). The intensities on the higher end of this range are sufficient to induce photoinhibition in many algae, which is when *P_max_* begins to decrease as light intensity increases (Kirk, 1994). The P-E curves for our species, however, do not show evidence of photoinhibition (Appendix S2). Cryptophytes are usually described as low light specialists (Gervais, 1997; Hoef-Emden & Archibald, 2016) but our data suggest that at least some cryptophytes photosynthesize relatively well at high light levels.

Photosynthetic rate often correlates with population growth rate (Falkowski et al., 1985; Coles & Jones, 2000) and is argued to represent the relative fitness of photosynthetic organisms (Violle et al., 2007). Therefore, species with the largest values for *P-E α* and *P_max_* are expected to have higher average fitness in across a wide range of light intensities, compared against species with lower trait values.

Prior work has shown that the relationship between maximum growth rate and the initial slope of a functional response is strongly positively correlated with body size (Kiørboe & Thomas, 2020). We accounted for this potential allometric relationship by including cell volume as a fixed effect in our PGLS models and did not see a significant correlation between it and the initial slope of a P-E curve or RLC. (Appendix S1: Table S3). Therefore, differences of cells’ volumes cannot explain the observed lack of a gleaner-opportunist trade-off.

There have been disagreements in the literature as to the relative importance, and even existence, of the gleaner-opportunist trade-off (Kiørboe & Thomas, 2020). It has long been assumed to play a crucial role in structuring ecological communities by allowing for coexistence between unequal competitors. We took a novel approach to evaluate the potential origin of this type of trade-off through evolutionary diversification of the traits themselves by using phylogenetic comparative methods to control for the evolutionary history of our focal species. As we did not find a gleaner-opportunist trade-off between photosynthetic traits our work supports the view that this trade-off may not be as widespread as previously assumed (Litchman et al., 2007; Isanta-Navarro et al., 2022; Yamamichi & Letten, 2022).

There are, however, caveats to our results. One is that we did not incorporate species-specific mortality rates into our experiment. With the inclusion of mortality rates, an opportunist is defined by the ratio of its maximum growth rate to its mortality rate rather than simply being the species with the higher maximum growth rate when a resource is abundant (Litchman & Klausmeier, 2001). The trade-off could then potentially emerge between a species with a low resource requirement and one with a high ratio of maximum growth rate to mortality rate (Litchman & Klausmeier, 2001). In the context of our study, the ratio for defining an opportunist would be the ratio of maximum photosynthetic rate to mortality rate. By ignoring mortality rates we are potentially overlooking a condition through which a gleaner-opportunist trade-off may manifest.

### No evidence of electron transport trade-offs

A possible explanation for the lack of a trade-off at the scale of overall photosynthesis is that an assimilation trade-off may exist in the electron transport chain of the light reactions but is masked by compensation in other parts of photosynthesis. This explanation is however excluded by our electron transport rate data, which showed no relationship between the initial slope of a species’ RLC and its maximum relative electron transport rate (Figure 3b).

The positive relationship between *P-E α* and *P_max_* could have arisen due to a mechanistic link between photosynthesis and electron transport. We expected species that were transferring more electrons between photosystems II and I to show a higher maximum photosynthetic rate as a higher rate of electron transport which would allow for more carbon to be fixed during the Calvin Cycle (Falkowski & Raven, 2007). We tested whether maximum photosynthetic rate was positively correlated with maximum rate of electron transfer. Somewhat surprisingly, we found no evidence of a relationship (Figure 3c), indicating that greater electron transport between photosystems does not yield greater photosynthesis. This relationship, however, would really be expected only if all the energy generated by electron transport is being used to fix carbon. Physiological plasticity in diverting energy generated through electron transport to alternative metabolic pathways may explain the absence of this correlation (Halsey & Jones, 2015).

### An escape from photosynthetic trade-offs?

Trade-offs promote ecological diversity by allowing competing species to coexist. Competitors may experience trade-offs in resource acquisition, resource allocation, differential predation, or dispersal ability; all these mechanisms can create the conditions necessary for stable coexistence (Chesson, 2000; Chase & Leibold, 2003; Ellner et al., 2019). Our data shows no evidence for a gleaner-opportunist trade-off between photosynthetic traits in cryptophytes. Thus, some cryptophytes should be strong competitors across a wide range of light environments. This lack of a trade-off, specifically between *P_max_* and *P-E α*, has been observed before and physiological mechanisms have been suggested as the cause of this relationship (Behrenfeld et al., 2008; Halsey et al., 2010).

In ecological comparisons of phytoplankton growing in different nutrient environments, Halsey et al. (2011; 2013; 2014) suggested positive covariation between *P_max_* and *P-E α* is driven by carbon metabolism occurring via different pathways. These researchers manipulated nitrogen or light to limit growth rates in green algae and diatoms, thereby producing positive covariation of *P_max_* and *P-E α* across environments due to changes in carbon metabolism. They argue that at low growth rates, induced by nitrogen or light limitation, a fixed transient carbon pool is quickly used for synthesis of ATP or NADPH, nucleic acids, and lipids. In contrast, at high growth rates fixed transient carbon is mostly stored as polysaccharides and used over a longer timescale. Thus, these within-species environmental manipulations recovered the same pattern as our across-species evaluation of evolutionary diversification, but it is unclear whether the proposed mechanism could be the same. First, in our study, all cultures were grown in nutrient-rich media for short periods of time, and thus none should have been nitrogen-limited. This is particularly true because our measurements were taken at mid-exponential phase. Second, we do not have information on the transient carbon pool or polysaccharide storage to characterize carbon metabolism in cryptophytes. Knowing a positive relationship exists, acquiring these types of data becomes a priority for future work.

The carbon metabolism hypothesis provides a potential explanation for the observed lack of gleaner-opportunist and power-efficiency trade-offs but does not rule out the possibility that trade-offs occur between other resource acquisition traits that we did not investigate. In fact, its empirical link to nitrogen limitation points to the importance of considering different types of resources. Trade-offs among resource acquisition traits in phytoplankton are well known, such as being a strong competitor for light but a weak competitor for nutrients (Tilman, 1977; Litchman & Klausmeier, 2008). Phytoplankton resource acquisition traits are remarkably plastic (Stomp et al., 2008; Hattich et al., 2016) so environmental variability such as intermittent predation pressure, sporadic nutrient limitation, interspecific competition, or temperature shifts could potentially drive the emergence of photosynthetic trade-offs. Over evolutionary timescales, cryptophytes that display low trait values for both *P_max_* and *P-E α* may have optimized other aspects of fitness to be excellent competitors for other resources, leading to trade-offs among traits not investigated in this study.

### Implications for natural phytoplankton communities

While our study aimed to broadly examine photosynthetic trade-offs through the gleaner-opportunist framework, our results also imply that the niches of cryptophytes in natural phytoplankton communities may be misunderstood. In particular, the realized niche of cryptophytes may have historically been interpreted as their fundamental niche. Cryptophytes are often described as low-light specialists (Hoef-Emden & Archibald, 2016) and frequently occur at deeper depths where less light is available (Gervais, 1997). At these depths, light attenuation leads to a decrease in the number of photons available for photosynthesis and alters the color of available light. Absorption by chlorophyll *a* and colored dissolved organic matter reduces the blue light present in the environment while green and red light are absorbed by accessory pigments like phycoerythrin and phycocyanin. In conjunction with the MAPPS data (Appendix S1: Figure S2), our results suggest that cryptophytes are not exceptional at capturing photons at low light intensities when compared to other taxa of phytoplankton, in contrast to expectations for low-light specialists. Furthermore, our P-E curves (Appendix S2) provide no evidence for photoinhibition at high light intensities. Cryptophytes instead have a broad fundamental niche with respect to light intensity and are better thought of as low-light tolerant rather than specialists. Cryptophytes may use their unique pigmentation to absorb colors of light that are not absorbed by taxa present at the surface. They could be exploiting fine gradations of light color rather than light intensity to carve out realized niches in different aquatic ecosystems. In addition to being able to exploit different colors of light, cryptophytes may succeed at deeper depths because due to greater nutrient availability than surface waters. Cryptophytes require nitrogen for phycobiliprotein synthesis (Doust et al., 2006) and have been shown to rapidly degrade these nitrogen-containing pigments when nitrogen is scarce (Da Silva et al., 2009). Living deeper may provide cryptophytes with access to sufficient nitrogen to synthesize phycobiliproteins, which then allows them to exploit wavelengths of light for photosynthesis that are not absorbed by other taxa.

### Future directions for trade-off research

Trade-offs may be dependent on the scale at which they are investigated (i.e., at the level of phenotype, genotype, population, or species) (Agrawal, 2020). For example, a relationship may be seen from a within-species comparison, but the same relationship may not manifest at the among-species level. This pattern of trade-offs occurring across different scales can be seen in trade-offs between milkweed leaf traits (Agrawal, 2020). A trade-off between leaf mass per area and cardenolide concentration can be seen between populations but is not apparent at the level of genotype or species (Agrawal, 2020). For plant traits, the general expectation is that trade-offs or predicted trade-offs will not be persistent across different scales (Agrawal, 2020). This paradigm of scale-dependent trade-offs may also be applicable to phytoplankton and may provide an investigative framework for future work on photosynthetic trait trade-offs.

Our study tested for photosynthetic trade-offs within an explicit resource exploitation framework while controlling for evolutionary history via phylogenetic comparative methods. This approach should be applicable for researchers working in systems where competition for light among photosynthetic organisms plays a strong role in structuring communities. This may include aquatic macrophytes (Sand-Jensen et al., 2007), forest communities (Onoda et al., 2015), grassland communities (Dybzinski & Tilman, 2007; Hautier et al., 2009), and coral endosymbionts (McIlroy et al., 2019). Examining strategies for light capture through pre-existing ecological frameworks can provide new perspectives and insights on how photosynthetic organisms interact with light in their environment.

## Acknowledgments

We thank Kristin Heidenreich, Krista Harmon, Cameron Riddick, and Patrick Lawson for help with culture maintenance and Jay Pinckney for the inspiration and equipment for measuring electron transport rate via rapid light curves. This study was supported by the National Science Foundation (NSF) Dimensions of Biodiversity program under grant #1542555 to T.L. Richardson and J. L. Dudycha.

## Author Contributions

The experiment was designed by JAS, TLR, and JLD. Data for the P-E curves and RLCs were collected by JAS. DNA extractions and phylogeny creation was done by MJG. Statistical analyses were done by JAS. The manuscript was written by JAS, MJG, and JLD with all authors contributing to the editing process.

