## Supplemental Appendix 1 for "A test of the gleaner-opportunist trade-off among photosynthetic traits in Cryptophyte algae"

Table of Contents

Section S1: Methods pgs. 2-6

Phytoplankton culture conditions pg. 2

Photosynthesis vs. irradiance curves pgs. 2-4

Rapid light curves- pgs. 4-5

UCE processing and phylogenetic inference pgs. 5-6

Sections S2: Results pgs.6-7

Photosynthesis and electron transport parameter estimates pgs. 6-7

Section S3: Tables and Figures pgs. 8-12

Table S1 pg. 8

Table S2 pg. 9

Table S3 pg. 10

Figure S1 pg. 11

Figure S2 pg. 12

References pgs. 13-16

Section S1: Methods

*Phytoplankton culture conditions*

Cultures were grown in nutrient replete medium at 17.5 °C (Table S1) with 30 μmol photons m^-2^ s^-1^ full spectrum irradiance on a 12:12 day:night cycle in a controlled environment chamber. Cultures were grown in 250 mL Pyrex bottles filled to 150 mL and repositioned daily, randomly, within the chamber. Neutral density filters were used to attenuate the light to 30 μmol photons m^-2^ s^-1^ where necessary. The light spectrum was measured with a StellarNet Blue Wave Mini Spectrometer (StellarNet, Tampa, FL, USA) (Figure S1). Cultures acclimated to growth conditions for 10 generations (~ 2-3 weeks) before measurements were made. During acclimation, cultures were transferred into new bottles once they were in mid-exponential growth phase. For experimental measurements, samples were collected only while cultures were in mid-exponential growth phase.

*Photosynthesis vs. irradiance (P-E) curves*

For each species, NaH^14^CO_3_ was added to each sample to achieve a final activity of approximately 3 μCi mL ^-1^. One mL of sample was dispensed into each of 16 scintillation vials. To obtain the total activity of ^14^C, three total count samples of 50 μL were dispensed into vials containing 250 μL of monoethanolamine and 5 mL of scintillation cocktail (EcoLumeTM; MP Biomedicals, Solon, OH, USA). Samples were exposed to light intensities ranging from 0 -1400 μmol photons m^-2^ s^-1^ and irradiance was measured with a quantum scalar irradiance sensor (Biospherical Instruments, Inc., San Diego, CA, USA) inserted into an empty scintillation vial. Samples were incubated with ^14^C for 20 minutes. Temperature during incubation was kept constant with a circulating water bath at 17.5 °C.

Chlorophyll-specific rates of photosynthesis were plotted against light intensity and data were fit with the equation of Platt et al., 1980 (Equation S1):

$\text{P}^{\text{B}}\text{= }\text{P}_{\text{S}}^{\text{B}}\left( \text{1-}\text{exp}^{-\left( \frac{\alpha\text{* E}}{\text{P}_{\text{S}}^{\text{B}}} \right)} \right)\left( \text{exp}^{-\left( \text{ }\frac{\text{β }\text{* }\text{E}}{\text{P}_{\text{S}}^{\text{B}}} \right)} \right)\text{ }$(Equation S1)

Where *P^B^* is the rate of photosynthesis scaled to chlorophyll *a* (g C *(g chl *a*)^-1^ h^-1^), $\text{P}_{\text{S}}^{\text{B}}$ is the maximum photosynthetic rate without photoinhibition (g C *(g chl *a*)^-1^ h^-1^), *E* is the light intensity (μmol photons m^-2^s^-1^), $\alpha$ is the initial slope of the P-E curve, (g C *(g chl *a*^-1^) h^-1^ (μmol photons m^-2^s^-1^)^-1^), and $\text{β }$(g C *(g chl *a*)^-1^ h^-1^) (μmol photons m^-2^s^-1^)^-1^) is the slope of the curve at supersaturating light intensities, when applicable. Curve fits were used to estimate $\alpha$, $\text{β}$, and $\text{P}_{\text{S}}^{\text{B}}$ . All fits were done with R version 3.6.2 using a non-linear least squares method via the “nlsLM” function in the package “minpack.lm” (Elzhov et al. 2016) The maximum rate of photosynthesis*, P_max_* (g C *(g chl *a*)^-1^ h^-1^), was calculated following Platt et al. 1980:

$P_{\max}= P_{S}^{B}\left( \frac{\alpha}{\alpha+\beta} \right)\left( \frac{\beta}{\alpha+\beta} \right)^{\frac{\beta}{\alpha}}$(Equation S2)

Total dissolved inorganic carbon (DIC) in whole samples for each species was determined prior to the addition of radioisotope with a LI-COR 820 CO_2_ Gas Analyzer (LI-COR, Lincoln, Nebraska, USA). Gaseous CO_2_ was evolved via the addition of H_2_SO_4_ to whole samples and the total DIC in the resultant gas was determined with an infrared optical sensor.

The chlorophyll *a* concentration of each species was measured to scale measurements of photosynthesis rate to a biomass index. A 5 mL sample of phytoplankton from each culture used for P-E curve measurements was filtered onto a GF/C filter and extracted overnight in the freezer in 5 mL of 90% acetone. The following day the chlorophyll concentration of the eluent was measured using a Turner Designs 10-Au fluorometer (Turner Designs, San Jose, CA, USA).

*Rapid light curves and electron transport activity*

Parameters of electron transport for each species were assessed via the generation of RLCs using pulse-amplitude-modulated (PAM) chlorophyll *a* fluorescence with a Walz Water-PAM (PAM, Heinz-Walz, Germany). Samples were dark-adapted for 20 minutes and then exposed to nine pulses of actinic light increasing in intensity with a range from 30 to 1300 μmol photons m^-2^s^-1^ with a 30 second interval between each light pulse. Curves were fit like Eq.1, with $P_{S}^{B}$ being replaced by *rETR_mPot_* and *P^B^* being replaced by *rETR* which yields:

$\text{rETR}\text{= }\mathrm{rETR}_{\mathrm{mPot}}\left( \text{1-}\text{exp}^{-\left( \frac{\alpha* E}{\mathrm{rETR}_{\mathrm{mPot}}} \right)} \right)\text{ }\left( \text{exp}^{-\left( \frac{\beta* E}{\mathrm{rETR}_{\mathrm{mPot}}} \right)} \right)\text{ }$(Equation S3)

Where *rETR* (μmol electrons m^-2^s^-1^) is the relative electron transport rate*, rETR_mPot_* (μmol electrons m^-2^s^-1^) is the maximum potential relative electron transport rate, $\alpha$ (μmol electrons/photon) is the initial slope of a RLC, and $\text{β}$ (μmol electrons/photon) is slope of the curve at supersaturating light intensities, and *E* is light intensity in units of μmol photons m^-2^s^-1^. This generated estimates of *RLC* $\alpha$, *rETR_mPot_*, and $\text{β}$ for each species. Estimates of *rETR_max_* (μmol electrons m^-2^s^-1^) were calculated following:

$\mathrm{rETR}_{\max}= \mathrm{rETR}_{\mathrm{mPot}}\left( \frac{\alpha}{\alpha+\beta} \right)\left( \frac{\beta}{\alpha+\beta} \right)^{\frac{\beta}{\alpha}}$(Equation S4)

Electron transport traits were measured in triplicate for each species. Note that only 14 species were used in the measurement of RLCs as our culture of one strain (*Cryptomonas sp*. CPCC 336) died before measurements could be taken.

*UCE processing and phylogenetic inference*

We downloaded the nuclear genomes of *Guillardia theta* CCMP3327 and *Baffinella frigidus* CCMP2293 from the Department of Energy (DOE) Joint Genome Institute (JGI) genome portal and used them as reference genomes for ultraconserved element (UCE) probe design (Curtis et al. 2012; Nordberg et al. 2014). We used Phyluce version 1.4 to identify, extract and validate 7,112 nucleotide probe sequences from 1,868 conserved nuclear genome loci (Faircloth et al. 2012; Faircloth 2016).

We extracted DNA using a standard cetyltrimethyl ammonium bromide (CTAB) method following Cunningham et al. (2019). DNA samples were checked for purity and integrity using a Nanodrop and agarose gel, respectively. RAPiD Genomics, LLC (Gainesville, Florida) synthesized 7,112 biotinylated 120-mer probes using our probe sequences. We sent DNA samples to RAPiD Genomics, LLC where target enrichment sequencing was performed using Illumina 2 X 150 bp reads.

We processed UCE with Phyluce version 1.5.0 (Faircloth et al. 2012; Faircloth 2016). Adapter and quality trimming of the raw paired end reads were carried out using the program illumiprocessor (Faircloth 2013; Bolger et al. 2014). We assembled contigs using a variety of *de novo* assemblers (Trinity, Velvet, ABySS) and kmer values (25-65) to accommodate differences in target enrichment for each cryptophyte strain (Zerbino and Birney 2008; Simpson et al. 2009; Grabherr et al. 2011). We selected the assemblies with the highest number of matched UCE probes based on the phyluce_assembly_match_contigs_to_probes script (Table S2). We then aligned contigs using MAFFT, resulting in an alignment matrix of 1,275,375 nucleotides across 1,515 UCE loci (Katoh et al. 2002). We only retained UCE loci that were found among at least 5 of the 16 taxa (35%) which resulted in 1,101 total UCE loci. RAxML version 8.0.19 was used for phylogenetic inference (Stamatakis 2014). We carried out the rapid hill-climbing mode using the autoMRE bootstrap search. The search terminated after 100 bootstrap inferences. These inferences were then mapped on to the best-scoring maximum likelihood tree resulting from 500 thorough replicates. We employed a general time reversal substitution model (GTR) and a gamma distribution. We transformed the RAxML phylogeny to a time proportional phylogeny with TreePL by specifying the age of the root as 1.0 (Smith and O’Meara 2012).

Section S2: Results

*Photosynthesis and electron transport parameter estimates*

Our estimates for *P_max_* (0.071-7.10 μgC (μgChl *a*)^-1^h^-1^) and *P-E* $\alpha$ (0.0035-0.11 (μgC (μgChl *a*)^-1^ h^-1^ (μmol photons m^-2^ s^-1^)^-1^)) trend towards the lower end of published estimates in the literature. An analysis of the Marine Primary Production: Model Parameters from Space (MAPPS) project database, which contains estimates of photosynthetic parameters from a global set of 5,711 P-E experiments with marine phytoplankton, found estimates for *P_max_* that range from 0.21-25.91 mgC (mgChl *a*)^-1^h^-1^ and estimates for *P-E* $\alpha$ that range from 0.002 to 0.373 mgC (mgChl *a*)^-1^ h^-1^ (μmol photons m^-2^ s^-1^)^-1^ (Bouman et al. 2018). Our measurements of *P_max_* and *P-E* $\alpha$ for cryptophytes are similar to the parameter estimates found by the MAPPS project; for the MAPPS data the mean for *P_max_* was 3.11 mgC (mgChl a)^-1^h^-1^ with a standard deviation of 2.28, while the mean for *P-E* $\alpha$ was 0.043 mgC (mgChl a)^-1^ h^-1^ (μmol photons m^-2^ s^-1^)^-1^ with a standard deviation of 0.034 (Bouman et al. 2018) (Fig. S2). In comparison, the mean for our *P_max_* estimates is 2.49 μgC (μChl a)^-1^ h^-1^with a standard deviation of 1.58, while the mean for our *P-E* $\alpha$ estimates is 0.044 μgC (μChl a)^-1^ h^-1^ (μmol photons m^-2^ s^-1^)^-1^ with a standard deviation of 0.030 (Fig. S2). Our data also does not suggest any photoinhibition occurring for the cryptophytes used in this study.

Our estimates for *P_max_* and *P-E* $\alpha$ for the 15 species of cryptophytes used in this study are consistent with the estimates from the MAPPS project, indicating that cryptophytes don’t have photosynthetic traits that fall outside the norm for phytoplankton. To the best of our knowledge, our data represents the most comprehensive estimates of cryptophyte photosynthetic traits. C^14^ incorporation-based estimates for *P_max_* and *P-E* $\alpha$ have been made for a strain of *Rhodomonas salina* CCAP 978/27 (Kaňa et al. 2012) and *Guillardia theta* (Cheregi et al. 2015) but do not appear to have been made for other strains of cryptophytes.

Few estimates of electron transport traits for cryptophytes have been reported, while reports for other taxa of phytoplankton are more common. *rETR_max_* and RLC $\alpha$ have been estimated for prymnesiophytes, chlorophytes, diatoms, dinoflagellates, and cyanobacteria (Wu and Song 2008; Gorai et al. 2014). Our estimates of *rETR_max_* and *RLC* $\alpha$ for the cryptophytes used in this study have a wider range for their parameter estimates but are otherwise comparable to estimates for other phytoplankton taxa. In terms of parameter estimates for cryptophytes *rETR_max_* was previously estimated for *Rhodomonas* *sp*. CCAC 0083 (Walter et al. 2017) and *rETR_max_*, normalized to number of cells, was estimated for *Geminigera* *cyrophila* (Koch and Trimborn 2019). As estimates of electron transport traits in cryptophytes remain scarce; our estimates of *rETR_max_* and *RLC* $\alpha$ represent a considerable addition to current estimates available in the literature.

Section S3: Tables and Figures

| Species | Culture Center ID | Habitat | Phycobiliprotein Absorption Peak | Cell Volume (μm^3^) | Light (μmol photons/m^2^s) | Temp (°C) | Growth Medium |
| --- | --- | --- | --- | --- | --- | --- | --- |
| *Chroomonas mesostigmatica* | CCMP 1168 | Marine | 645 | 134 | 30 | 20 | f/2-Si |
| *Chroomonas nordstedtii* | NIES 708 | Freshwater | 630 | 203 | 30 | 15 | MWC+Se |
| *Chroomonas placoidea* | NIES705 | Marine | 645 | 123 | 30 | 15 | L1-Si+ Organics |
| *Chroomonas sp.* | SCCAP K-1623 | Freshwater | 630 | 369 | 30 | 15 | MWC+Se |
| *Cryptomonas ovata* | UTEX 2783 | Freshwater | 566 | 1079 | 30 | 20 | MWC+Se |
| *Cryptomonas sp.* | CPCC 336 | Marine | 566 | 323 | 30 | 20 | Dy-V |
| *Goniomonas avonlea* | CCMP 3327 | Marine | - | 101 | 30 | 20 | FSW+ Organics+ Bacteria |
| *Guillardia theta* | CCMP 327 | Marine | 545 | 139 | 30 | 20 | H/2 |
| *Hemiselmis cryptochromatica* | CCMP 1181 | Marine | 569 | 30 | 30 | 15 | L1+NH4 |
| *Hemiselmis pacifica* | CCMP 706 | Marine | 577 | 91 | 30 | 15 | L1-Si |
| *Hemiselmis tepida* | CCMP 443 | Marine | 615 | 23 | 30 | 20 | L1+NH4 |
| *Proteomonas sulcata* | CCMP 1175 | Marine | 545 | 279 | 30 | 20 | f/2-Si |
| *Rhodomonas minuta* | CPCC 344 | Freshwater | 545 | 111 | 30 | 20 | MWC+Se |
| *Rhodomonas salina* | CCMP 1319 | Marine | 545 | 213 | 30 | 15 | L1-Si |
| *Teleaulax sp.* | RCC 4857 | Marine | 545 | 568 | 30 | 20 | f/2-Si |
| *Unid sp.** | ZIGY 001 | Freshwater | 566 | 399 | 30 | 20 | WH |

Table S1: List of species used in the study with species characteristics and stock culture conditions. As it is heterotrophic, *Goniomonas avonlea* does not have any phycobiliproteins.

*Strain was isolated from the field by the authors and is therefore not available in commercial culture collections

| Species | Culture Center ID | Assembler + kmer |
| --- | --- | --- |
| *Chroomonas mesostigmatica* | CCMP 1168 | Abyss k35 |
| *Chroomonas nordstedtii* | NIES 708 | Velvet k25 |
| *Chroomonas placoidea* | NIES 705 | Velvet k35 |
| *Chroomonas sp* | SCCAP K 1623 | Velvet k35 |
| *Cryptomonas ovata* | UTEX LB 2783 | Velvet k35 |
| *Cryptomonas sp* | CPCC 336 | Abyss k25 |
| *Goniomonas avonlea* | CCMP 3327 | Abyss k45 |
| *Guillardia theta* | CCMP 327 | Trinity N/A |
| *Hemiselmis cryptochromatica* | CCMP 1181 | Velvet k35 |
| *Hemiselmis pacifica* | CCMP 706 | Velvet k35 |
| *Hemiselmis tepida* | CCMP 443 | Velvet k35 |
| *Proteomonas sulcata* | CCMP 1175 | Velvet k35 |
| *Rhodomonas minuta* | CPCC 344 | Velvet k35 |
| *Rhodomonas salina* | CCMP 1319 | Velvet k35 |
| *Teleaulax sp* | RCC 4857 | Abyss k25 |
| Unid sp | Zigy 001 | Velvet k25 |

Table S2: Assembler used and matched number of UCE probes for each cryptophyte species

| PGLS Model Response Variable | Predictors | *F* | Df | Adjusted *R^2^* | *p* |
| --- | --- | --- | --- | --- | --- |
| P_max_ | Intercept | 3.195 | 4,10 | 0.3854 | 0.52 |
|  | Alpha |  |  |  | **0.012** |
|  | Cell Volume |  |  |  | 0.44 |
|  | PBP |  |  |  | 0.55 |
|  | Habitat |  |  |  | 0.86 |
| P_max_ | Intercept | 0.1968 | 4,9 | -0.3283 | 0.33 |
|  | ETR_max_ |  |  |  | 0.89 |
|  | Cell Volume |  |  |  | 0.73 |
|  | PBP |  |  |  | 0.44 |
|  | Habitat |  |  |  | 0.76 |
| ETR_max_ | Intercept | 1.286 | 3,10 | 0.06188 | 0.098 |
|  | Cell Volume |  |  |  | 0.65 |
|  | PBP |  |  |  | 0.17 |
|  | Habitat |  |  |  | 0.34 |

Table S3: Summary statistics for the full PGLS models that include cell volume, phycobiliprotein absorption peak, and habitat (marine or freshwater) as predictor variables. Degrees of freedom are indicated as subscripts to each *F* statistic and significant *p* values are in bold. The full model of ETRmax~ETRalpha +Marine+PBP+Cell Volume does not produce a result due to an optimization error, so the results are shown without ETR alpha as a predictor variable.

PAR (μmol photons m^-2^ s^-1^)

Wavelength (nm)

Figure S1: Light spectrum of incubator used to grow experimental cultures

|  |  |
| --- | --- |
| 400 | 0 |
| 400.5 | 0 |
| 401 | 0 |
| 401.5 | 0 |
| 402 | 0 |
| 402.5 | 0 |
| 403 | 0.0029516 |
| 403.5 | 0.014228 |
| 404 | 0.027876 |
| 404.5 | 0.035551 |
| 405 | 0.032226 |
| 405.5 | 0.01937 |
| 406 | 0.0033992 |
| 406.5 | 0 |
| 407 | 0 |
| 407.5 | 0 |
| 408 | 0 |
| 408.5 | 0 |
| 409 | 0 |
| 409.5 | 0 |
| 410 | 0 |
| 410.5 | 0 |
| 411 | 0 |
| 411.5 | 0 |
| 412 | 0 |
| 412.5 | 0 |
| 413 | 0 |
| 413.5 | 0 |
| 414 | 0 |
| 414.5 | 0 |
| 415 | 0 |
| 415.5 | 0 |
| 416 | 0 |
| 416.5 | 0 |
| 417 | 0 |
| 417.5 | 0 |
| 418 | 0 |
| 418.5 | 0 |
| 419 | 0 |
| 419.5 | 0.0003942 |
| 420 | 0.0039752 |
| 420.5 | 0.0064442 |
| 421 | 0.0070332 |
| 421.5 | 0.0061279 |
| 422 | 0.0030105 |
| 422.5 | 0.00041131 |
| 423 | 0 |
| 423.5 | 0.00012322 |
| 424 | 0.0022182 |
| 424.5 | 0.0042137 |
| 425 | 0.0039349 |
| 425.5 | 0.0035993 |
| 426 | 0.0023199 |
| 426.5 | 0.0022874 |
| 427 | 0.0043892 |
| 427.5 | 0.0075918 |
| 428 | 0.0090271 |
| 428.5 | 0.0081211 |
| 429 | 0.0069936 |
| 429.5 | 0.0053059 |
| 430 | 0.0053146 |
| 430.5 | 0.0096789 |
| 431 | 0.015014 |
| 431.5 | 0.019949 |
| 432 | 0.024341 |
| 432.5 | 0.025673 |
| 433 | 0.023835 |
| 433.5 | 0.030358 |
| 434 | 0.067773 |
| 434.5 | 0.14184 |
| 435 | 0.2285 |
| 435.5 | 0.2861 |
| 436 | 0.27279 |
| 436.5 | 0.20895 |
| 437 | 0.12722 |
| 437.5 | 0.066853 |
| 438 | 0.038924 |
| 438.5 | 0.028984 |
| 439 | 0.026897 |
| 439.5 | 0.028051 |
| 440 | 0.02683 |
| 440.5 | 0.025734 |
| 441 | 0.025239 |
| 441.5 | 0.023684 |
| 442 | 0.023492 |
| 442.5 | 0.021992 |
| 443 | 0.02069 |
| 443.5 | 0.021849 |
| 444 | 0.025031 |
| 444.5 | 0.027855 |
| 445 | 0.029429 |
| 445.5 | 0.028744 |
| 446 | 0.028031 |
| 446.5 | 0.026979 |
| 447 | 0.0253 |
| 447.5 | 0.024896 |
| 448 | 0.020976 |
| 448.5 | 0.017455 |
| 449 | 0.016376 |
| 449.5 | 0.01559 |
| 450 | 0.017536 |
| 450.5 | 0.020302 |
| 451 | 0.020103 |
| 451.5 | 0.018558 |
| 452 | 0.017544 |
| 452.5 | 0.018233 |
| 453 | 0.019175 |
| 453.5 | 0.019603 |
| 454 | 0.019953 |
| 454.5 | 0.019495 |
| 455 | 0.022239 |
| 455.5 | 0.024705 |
| 456 | 0.024918 |
| 456.5 | 0.024397 |
| 457 | 0.021233 |
| 457.5 | 0.01846 |
| 458 | 0.014959 |
| 458.5 | 0.010306 |
| 459 | 0.0068341 |
| 459.5 | 0.0054248 |
| 460 | 0.0078581 |
| 460.5 | 0.012251 |
| 461 | 0.013558 |
| 461.5 | 0.012947 |
| 462 | 0.01183 |
| 462.5 | 0.011818 |
| 463 | 0.01566 |
| 463.5 | 0.019137 |
| 464 | 0.018873 |
| 464.5 | 0.01491 |
| 465 | 0.010819 |
| 465.5 | 0.0077434 |
| 466 | 0.006166 |
| 466.5 | 0.006133 |
| 467 | 0.0054609 |
| 467.5 | 0.0044632 |
| 468 | 0.0072908 |
| 468.5 | 0.011668 |
| 469 | 0.01295 |
| 469.5 | 0.011432 |
| 470 | 0.0082727 |
| 470.5 | 0.0056005 |
| 471 | 0.0049853 |
| 471.5 | 0.0071991 |
| 472 | 0.0074794 |
| 472.5 | 0.0053521 |
| 473 | 0.0039952 |
| 473.5 | 0.0031478 |
| 474 | 0.0044063 |
| 474.5 | 0.0085969 |
| 475 | 0.011356 |
| 475.5 | 0.013179 |
| 476 | 0.013582 |
| 476.5 | 0.010405 |
| 477 | 0.0093753 |
| 477.5 | 0.0099353 |
| 478 | 0.010528 |
| 478.5 | 0.012769 |
| 479 | 0.013831 |
| 479.5 | 0.013953 |
| 480 | 0.014652 |
| 480.5 | 0.015505 |
| 481 | 0.017089 |
| 481.5 | 0.0186 |
| 482 | 0.021104 |
| 482.5 | 0.025689 |
| 483 | 0.030449 |
| 483.5 | 0.036142 |
| 484 | 0.043991 |
| 484.5 | 0.053086 |
| 485 | 0.063214 |
| 485.5 | 0.074241 |
| 486 | 0.083311 |
| 486.5 | 0.089473 |
| 487 | 0.09254 |
| 487.5 | 0.092779 |
| 488 | 0.091167 |
| 488.5 | 0.088243 |
| 489 | 0.085182 |
| 489.5 | 0.081671 |
| 490 | 0.077734 |
| 490.5 | 0.074212 |
| 491 | 0.071443 |
| 491.5 | 0.069561 |
| 492 | 0.066679 |
| 492.5 | 0.063988 |
| 493 | 0.061971 |
| 493.5 | 0.059882 |
| 494 | 0.058201 |
| 494.5 | 0.05551 |
| 495 | 0.05204 |
| 495.5 | 0.048716 |
| 496 | 0.045945 |
| 496.5 | 0.044156 |
| 497 | 0.041896 |
| 497.5 | 0.03851 |
| 498 | 0.035221 |
| 498.5 | 0.031515 |
| 499 | 0.028219 |
| 499.5 | 0.025575 |

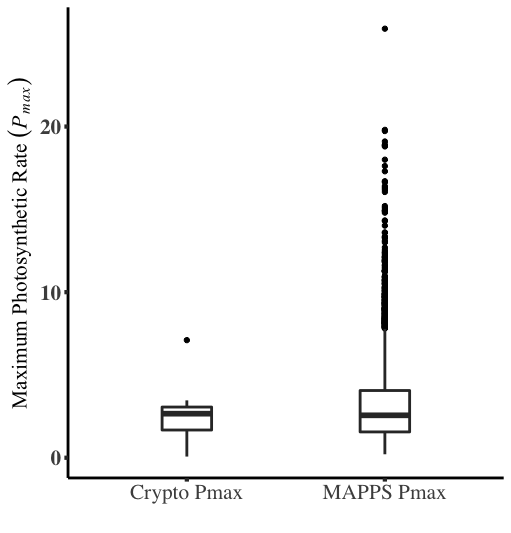

**(b)**

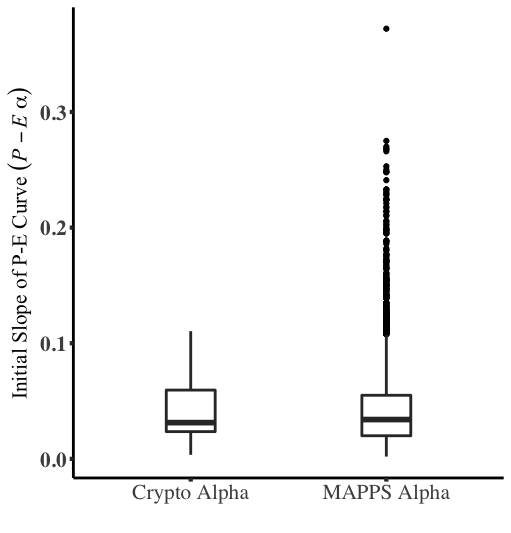

**(a)**

Figure S2: Boxplots showing comparisons of a) of the initial slope of a P-E curve, *P-E* $\alpha$, and b) comparison of maximum photosynthetic rate, *P_max_*, across our data and the MAPPS project data. Units for panel A are carbon per chlorophyll *a* per hour per irradiance as we measured *P-E* $\alpha$ in microgram C per microgram Chl *a* whereas the MAPPS data is in milligram C per milligram chlorophyll *a*. Units for panel B are carbon per chlorophyll *a* per hour where we measured *P_max_* in microgram C per microgram chlorophyll *a* and the MAPPS data is in milligram C per milligram chlorophyll *a*. The data collected from this studied is labeled Crypto Alpha and Crypto Pmax whereas the MAPPS data is labeled MAPPS Alpha and MAPPS Pmax. MAPPS data distributions were recreated using the data compiled by Bouman et al. 2018 and available as a supplemental file to that publication.

loci. *Bioinformatics, 32,* 786-788. https://doi.org/10.1093/bioinformatics/btv646

Faircloth B.C., McCormack, J.E., Crawford, N.G., Harvey, M.G., Brumfield, R.T., & Glenn,

T.C. (2012). Ultraconserved elements anchor thousands of genetic markers spanning multiple evolutionary timescales. *Systematic Biology, 61,*717-726. https://doi.org/10.1093/sysbio/sys004

Gorai, T., Katayama, T., Obata, M., Murata, A., & Taguchi, S. (2014). Low blue light enhances growth rate, light absorption, and photosynthetic characteristics of four marine phytoplankton species. *Journal of Experimental Marine Biology and Ecology,* *459*, 87–95. https://doi.org/10.1016/j.jembe.2014.05.013

Grabherr M.G., Haas, B.J., Yassour, M., Levin, J.Z., Thompson, D.A., Amit, I., Adiconis, X.,

Fan, L., Raychowdhury, R., Zeng, Q., Chen, Z., Mauceli, E., Hacohen, N., Gnirke, A., Rhind, N., di Palma, F., Birren, B.W., Nusbaum, C., Lindblad-Toh, K., … Regev, A. (2011). Trinity: reconstructing a full-length transcriptome without a genome from RNA-Seq data. *Nature Biotechnology, 29,* 644-652. https://dx.doi.org/10.1038/nbt.1883

Kaňa, R., Kotabová, E., Sobotka, R., & Prášil, O. (2012). Non-photochemical quenching in cryptophyte alga *Rhodomonas salina* is located in chlorophyll a/c antennae r. *PLoS ONE, 7,* e29700. https://doi.org/10.1371/journal.pone.0029700

Katoh K., Misawa, K., Kuma, K., & Miyata, T. (2002). MAFFT: a novel method for rapid

multiple sequence alignment based on fast Fourier transform. *Nucleic Acids Research, 30*, 3059-3066. https://doi.org/10.1093/nar/gkf436

Koch, F., & Trimborn, S. (2019). Limitation by Fe, Zn, Co, and B12 results in similar physiological responses in two antarctic phytoplankton species. *Froniers in Marine Science, 6*, 514. https://doi.org/10.3389/fmars.2019.00514

Nordberg H., Cantor, M., Dusheyko, S., Hua, S., Poliakov, A., Shabalov, I., Smirnova, T.,

Grigoriev, I., & Dubchak, I. (2014). The genome portal of the Department of Energy Joint Genome Institute: 2014 updates. *Nucleic Acids Research, 42*, D26-31.

https://doi.org/10.1093/nar/gkt1069

R Core Team 2020. R: A Language and Environment for Statistical Computing. R

Foundation for Statistical Computing, Vienna, Austria. https://www.R-project.org/.

Simpson, J.T., Wong, K., Jackman, S.D., Schein, J.E., Jones, S.J.M., & Birol, I. (2009). ABySS:

A parallel assembler for short read sequence data. *Genome Research, 19*, 1117-1123. https://doi.org/10.1101/gr.089532.108

Smith, S.A., & O’Meara, B.C. (2012). treePL: divergence time estimation using penalized l

ikelihood for large phylogenies. *Bioinformatics, 28*, 2689-2690. https://doi.org/10.1093/bioinformatics/bts492

Stamatakis, A. (2014). RAxML version 8: a tool for phylogenetic analysis and post-analysis of

large phylogenies. *Bioinformatics, 30*, 1312-1313. https://doi.org/10.1093/bioinformatics/btu033

Walter, B., Peters, J., & van Beusekom, J.E.E. (2017). The effect of constant darkness and short light periods on the survival and physiological fitness of two phytoplankton species and their growth potential after re-illumination. *Aquatic Ecology, 51*, 591–603. https://doi.org/10.1007/s10452-017-9638-z

York.

Wu, Z., & Song, L. (2008). Physiological comparison between colonial and unicellular forms of

*Microcystis aeruginosa* Kütz. (Cyanobacteria). *Phycologia, 47*, 98–104. https://doi.org/10.2216/07-49.1

Zerbino, D.R., & Birner, E. (2008). Velvet: algorithms for de novo short read assembly using de

graphs. *Genome Research, 18*, 821-829. https://doi.org/10.1101/gr.074492.107
