## Supplemental Appendix 2 for "A test of the gleaner-opportunist trade-off among photosynthetic traits in Cryptophyte algae"

P-E curves for all Cryptophyte species examined in our study. In order, curves are for a) *Chroomonas mesostigmatica*, b) *Chroomonas nordstedtii*, c) *Chroomonas placoidea*, d) *Chroomonas sp. K-1623*, e) *Cryptomonas ovata*, f) *Cryptomonas sp. 336*, g) *Guillardia theta*, h) *Hemiselmis cryptochromatica*, i) *Hemiselmis pacifica*, j) *Hemiselmis tepida,* k) *Proteomonas sulcate*, l) *Rhodomonas minuta*, m) *Rhodomonas salina*, n) *Teleaulax sp.4857*, o) *Unid. sp. ZIGY 001*

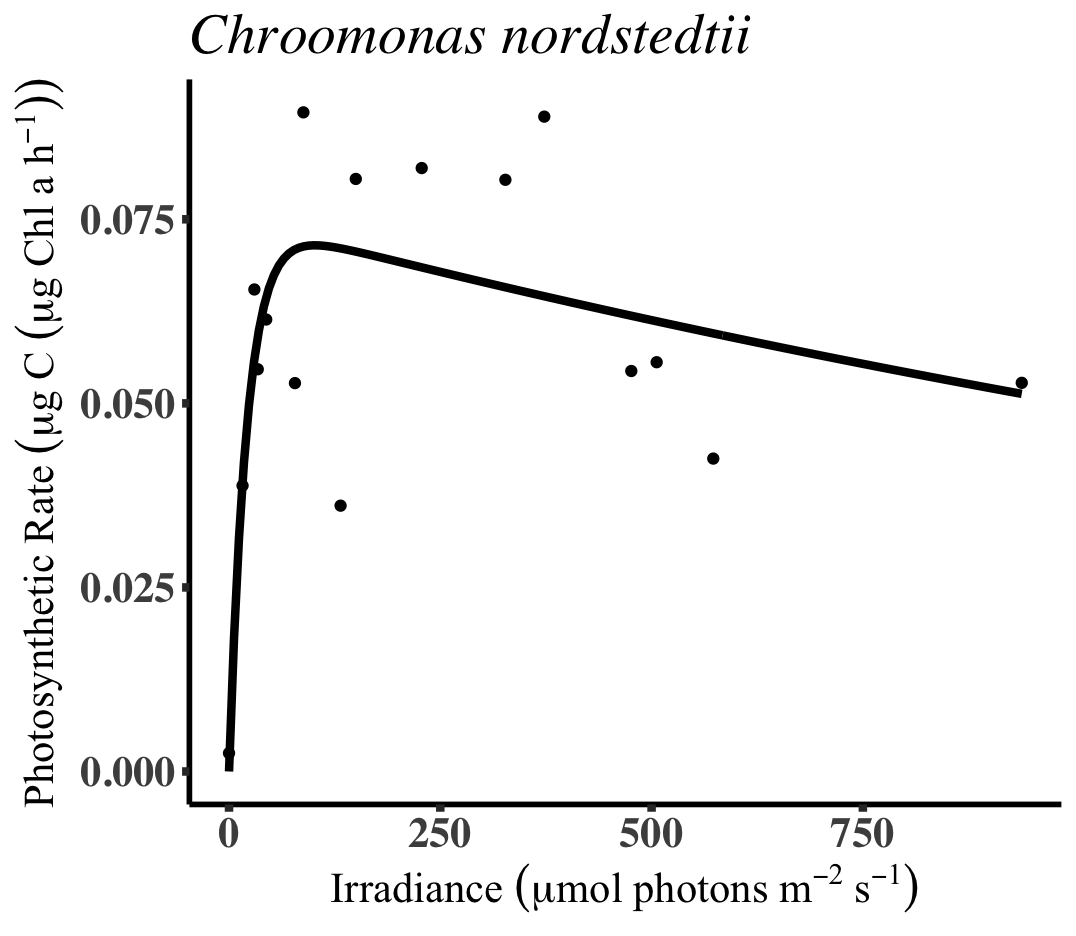

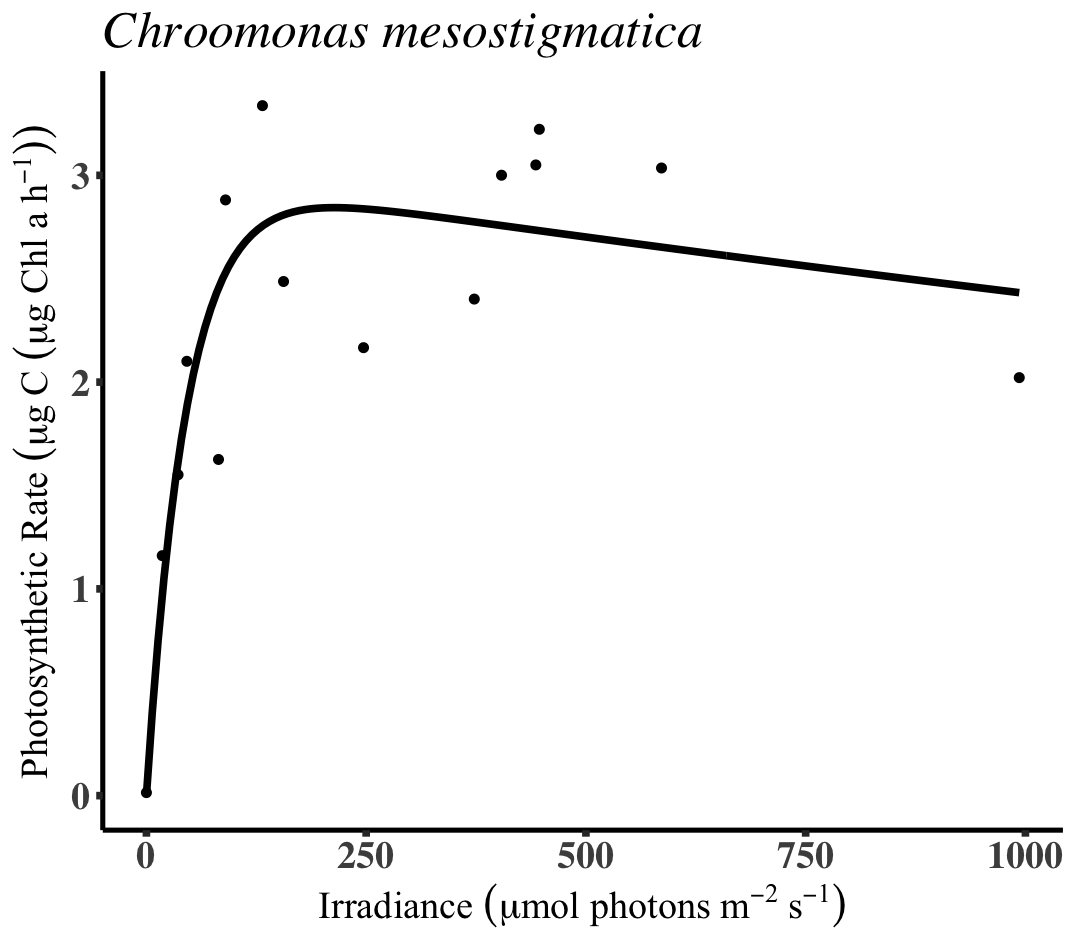

**b)**

**a)**

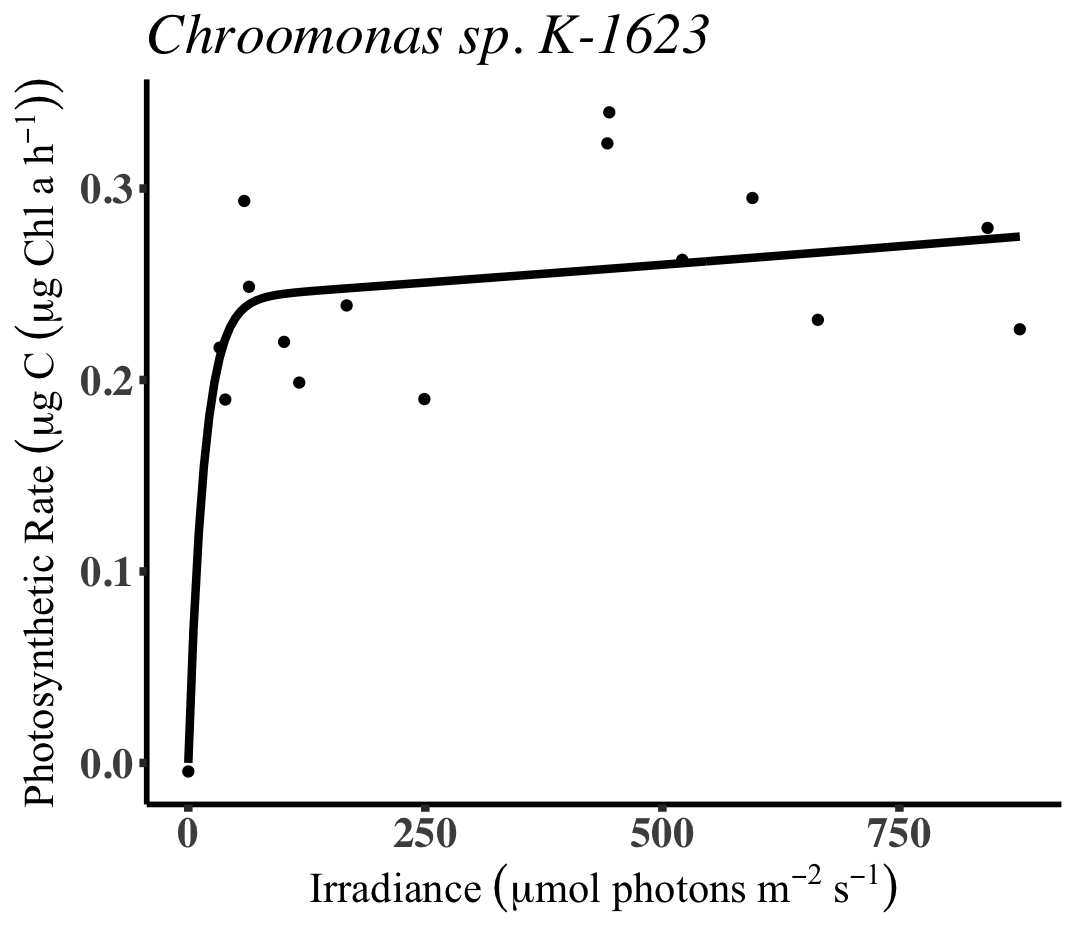

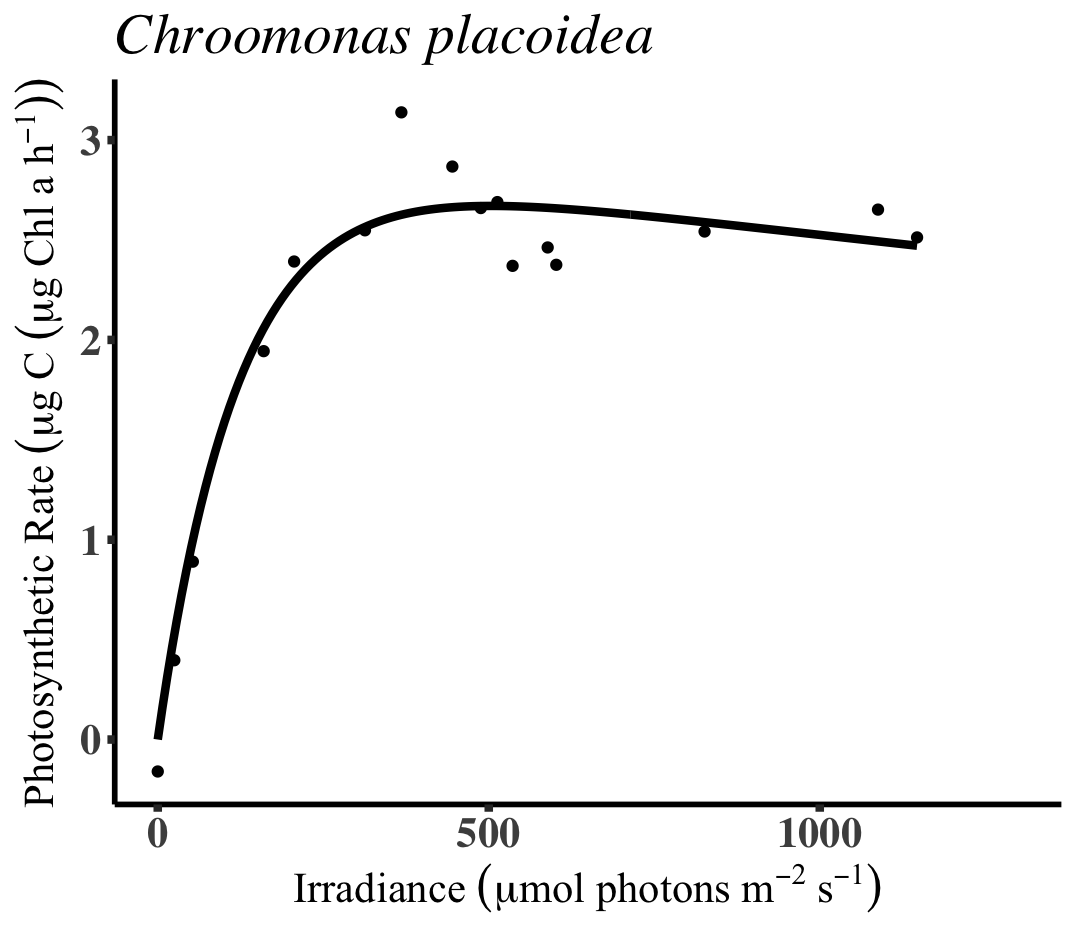

**d)**

**c)**

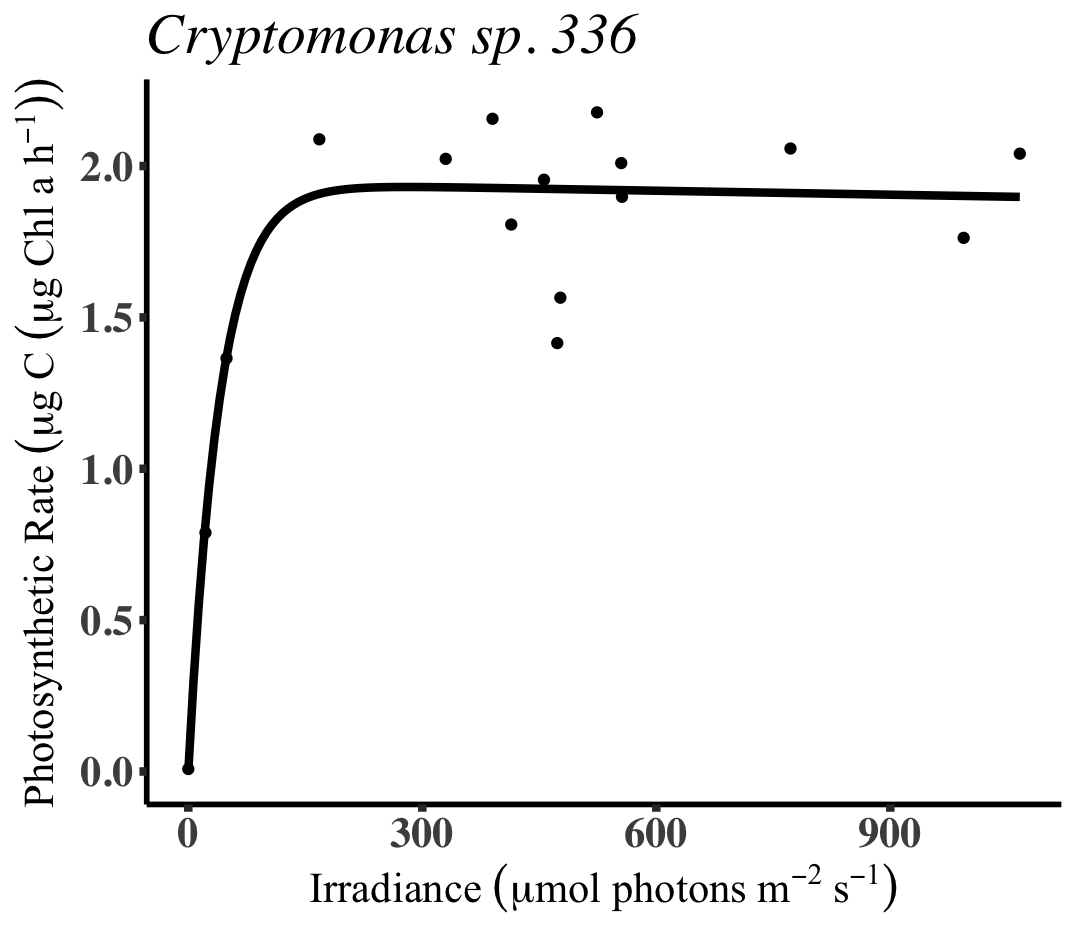

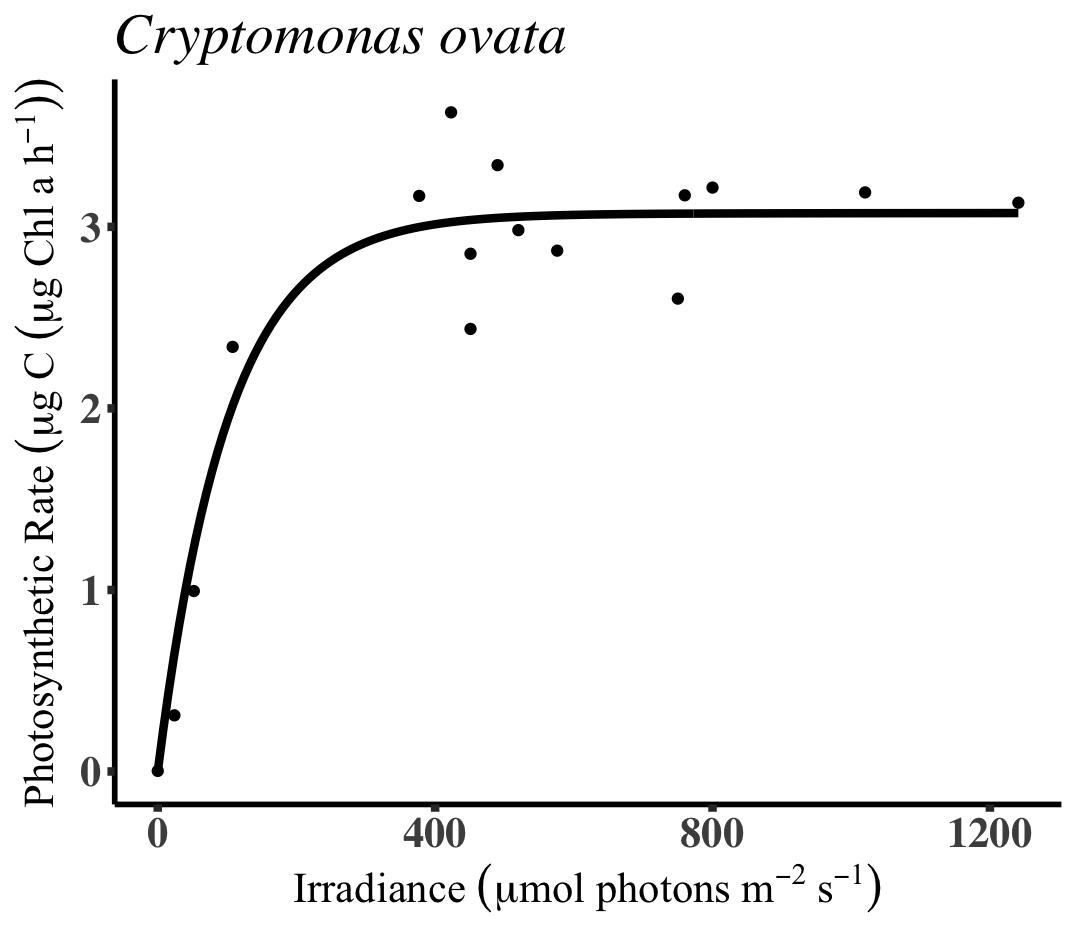

**f)**

**e)**

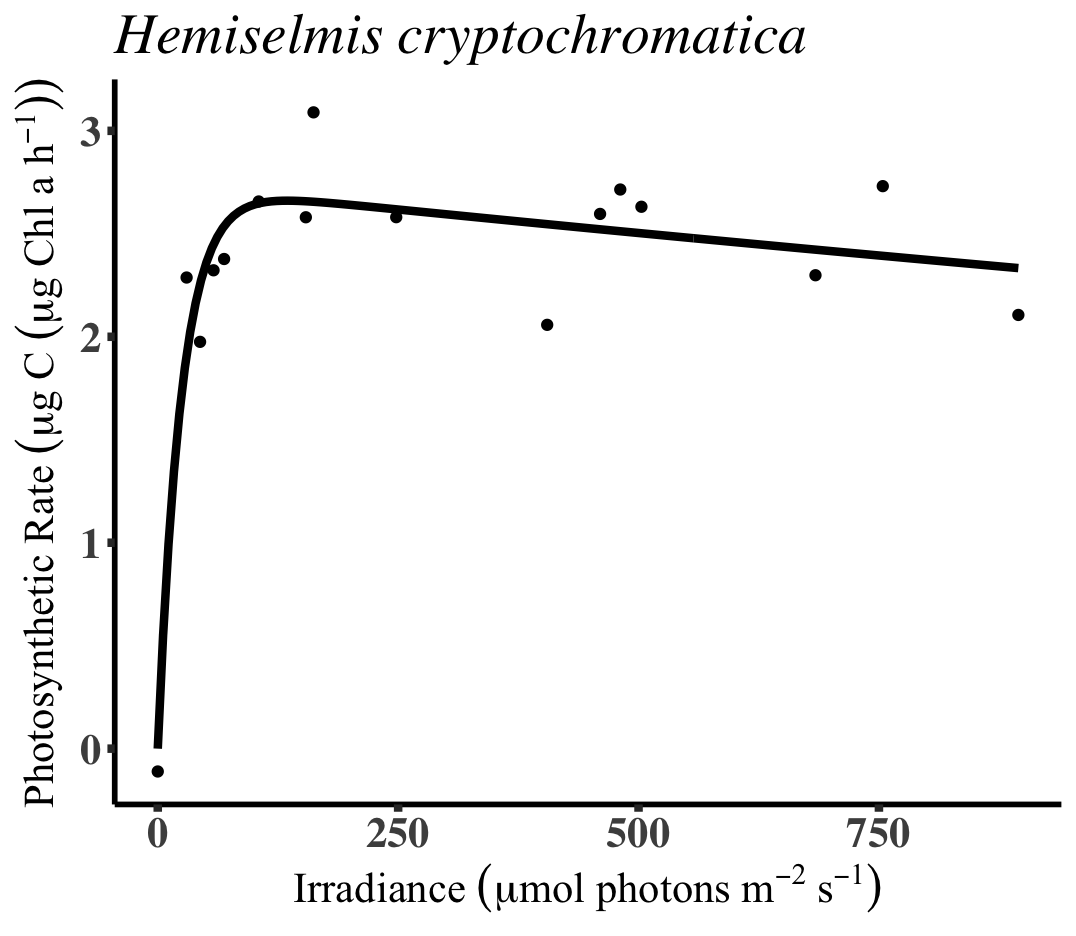

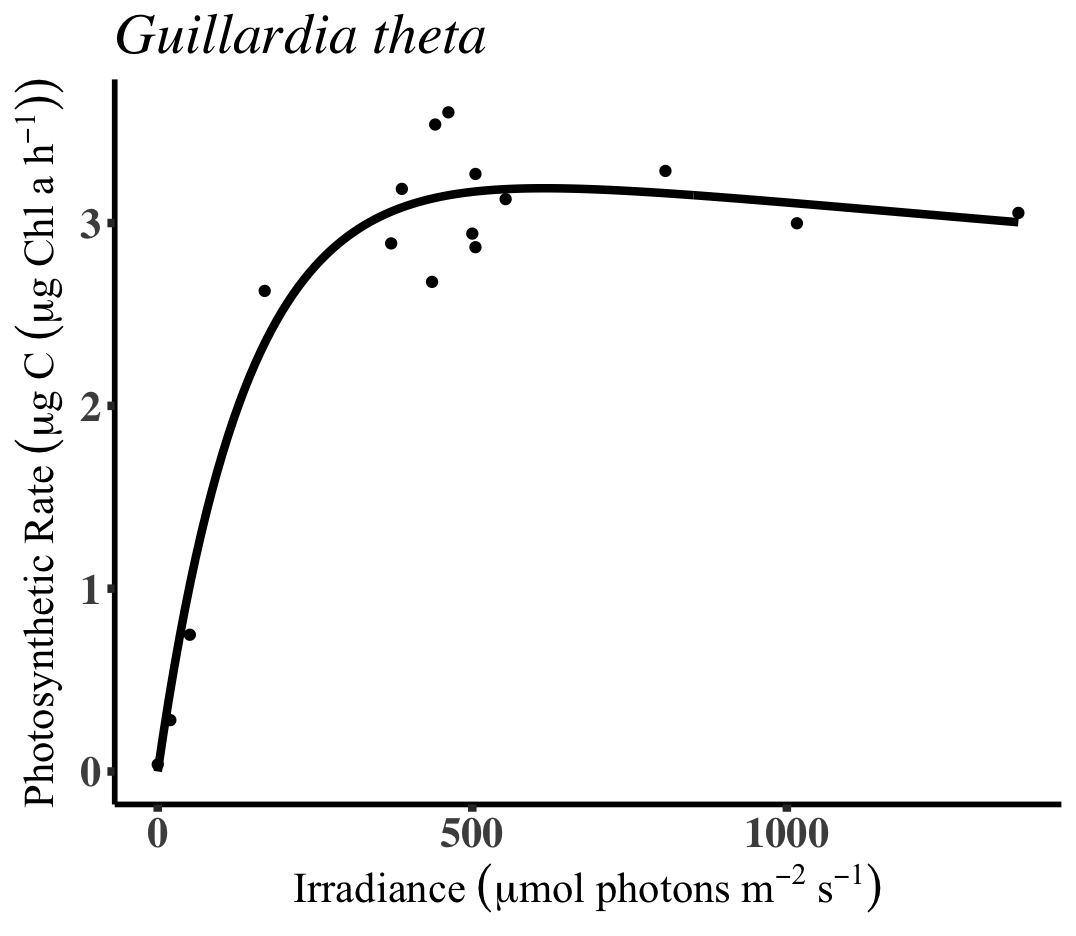

**h)**

**g)**

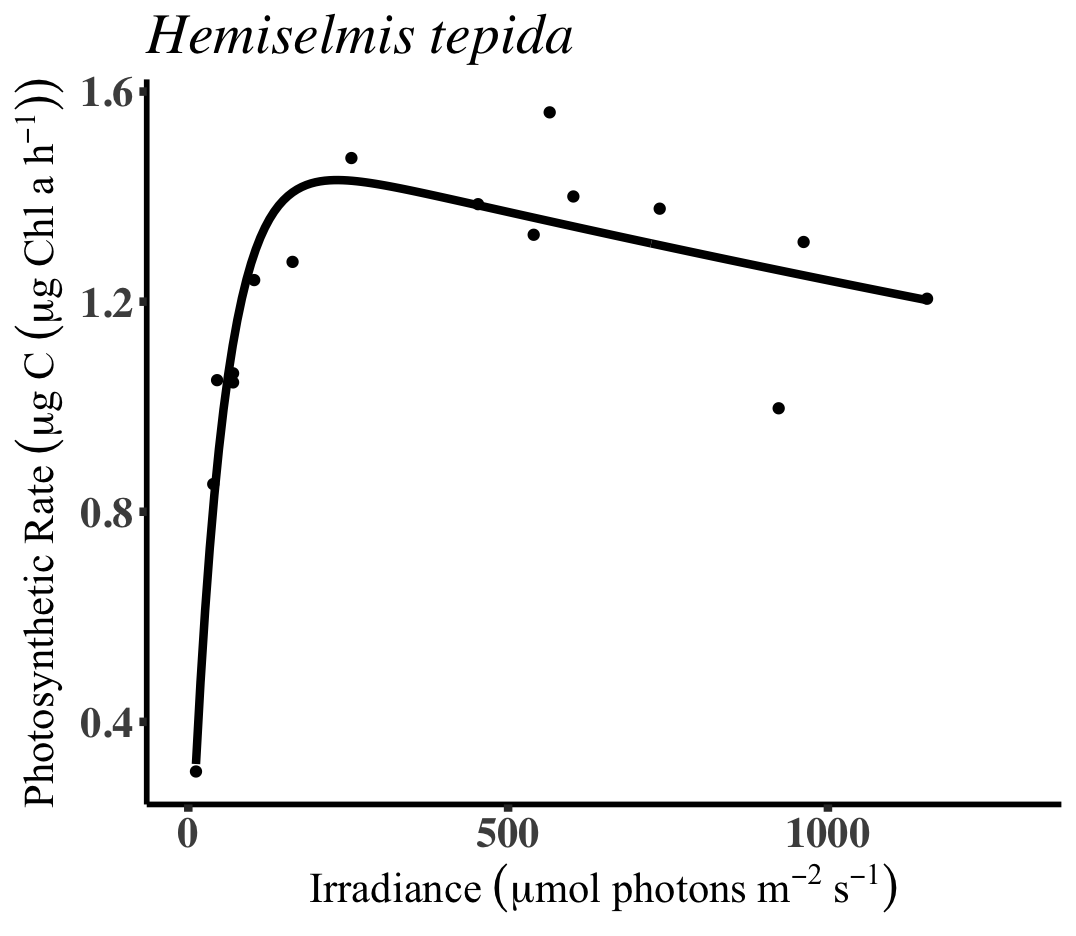

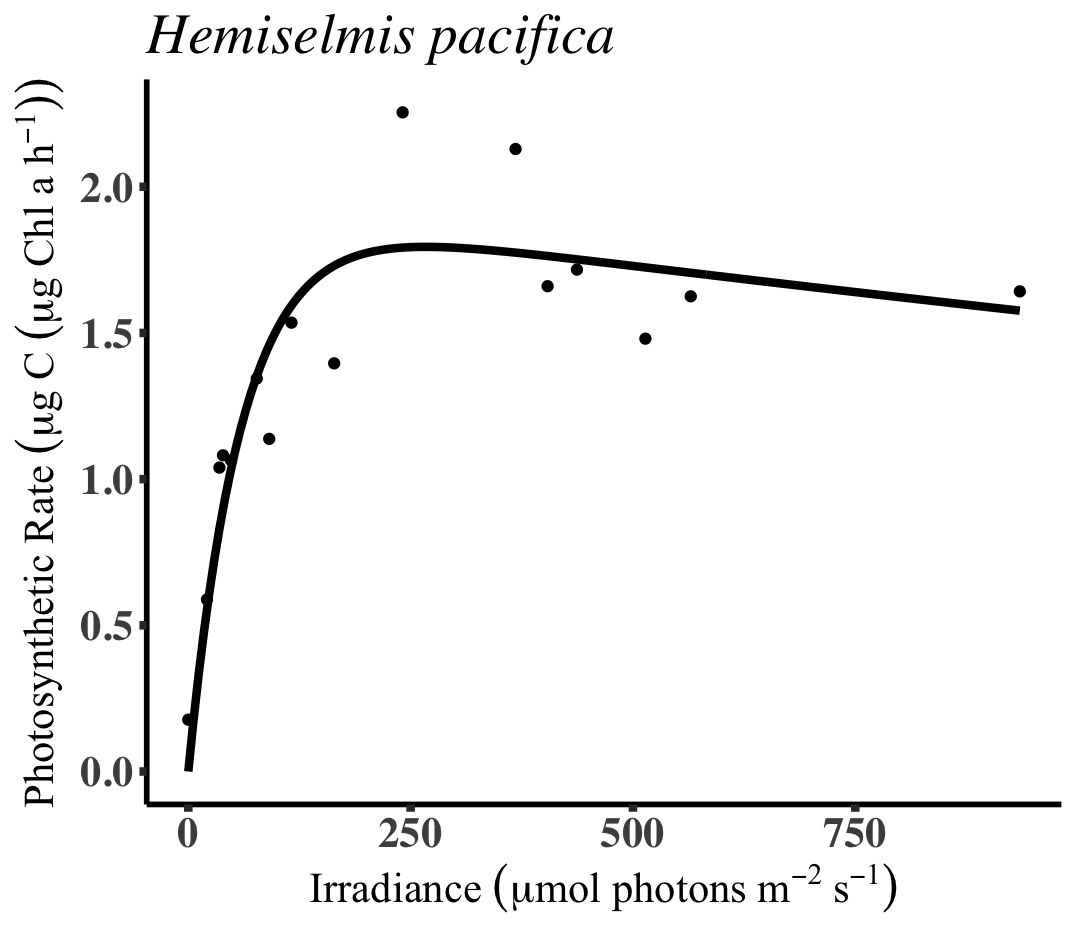

**j)**

**i)**

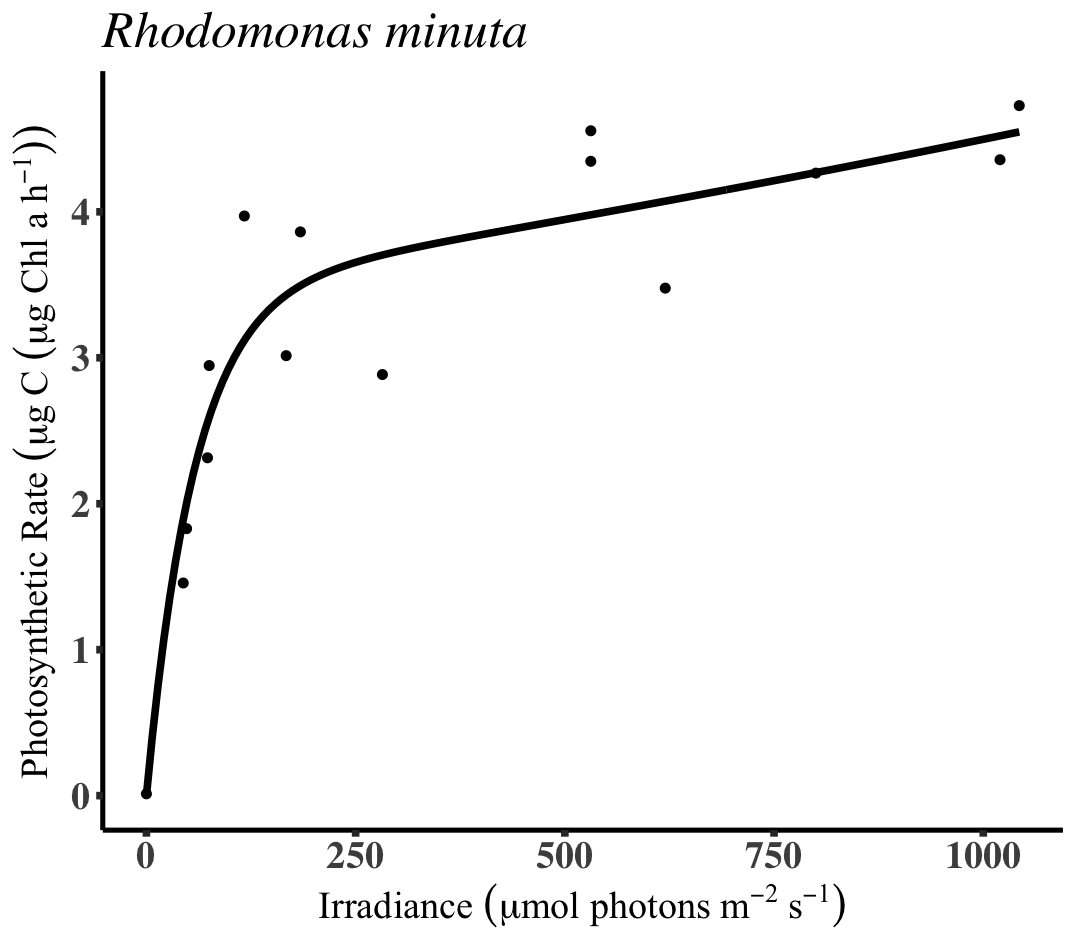

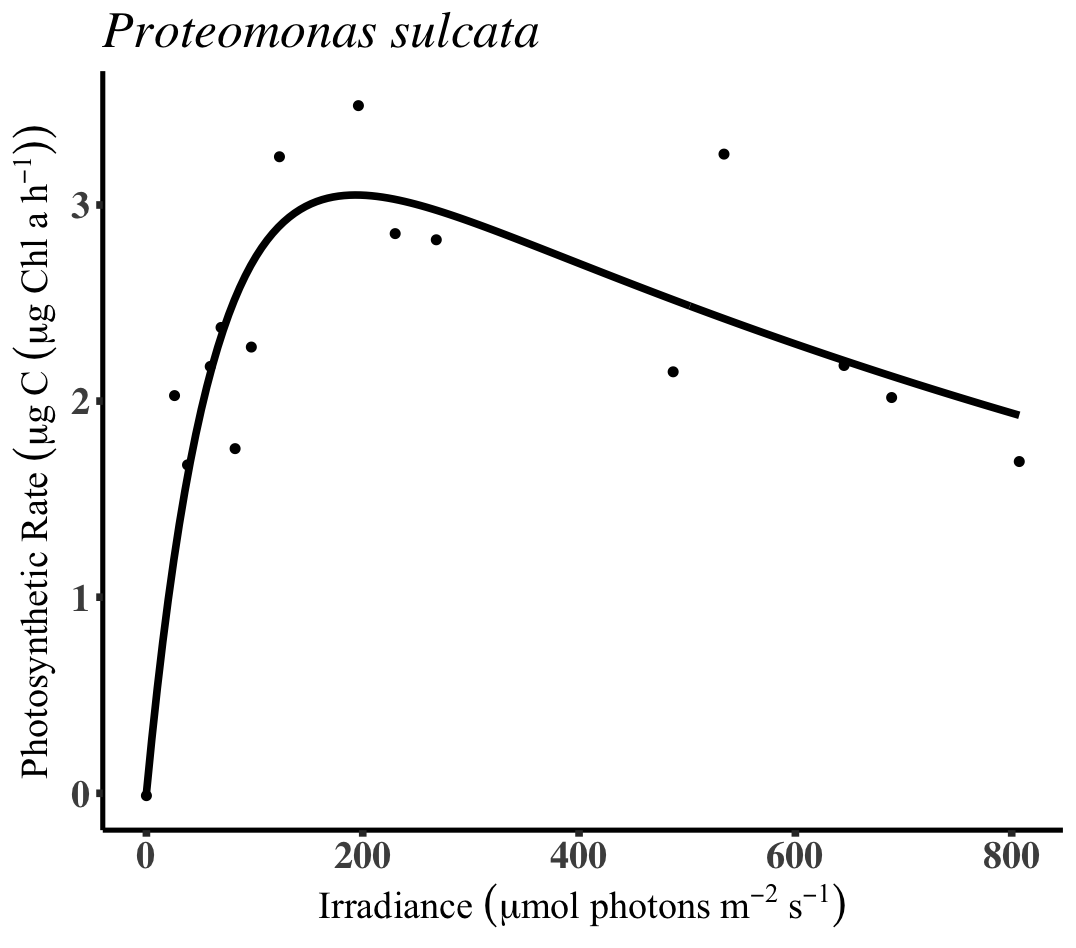

**l)**

**k)**

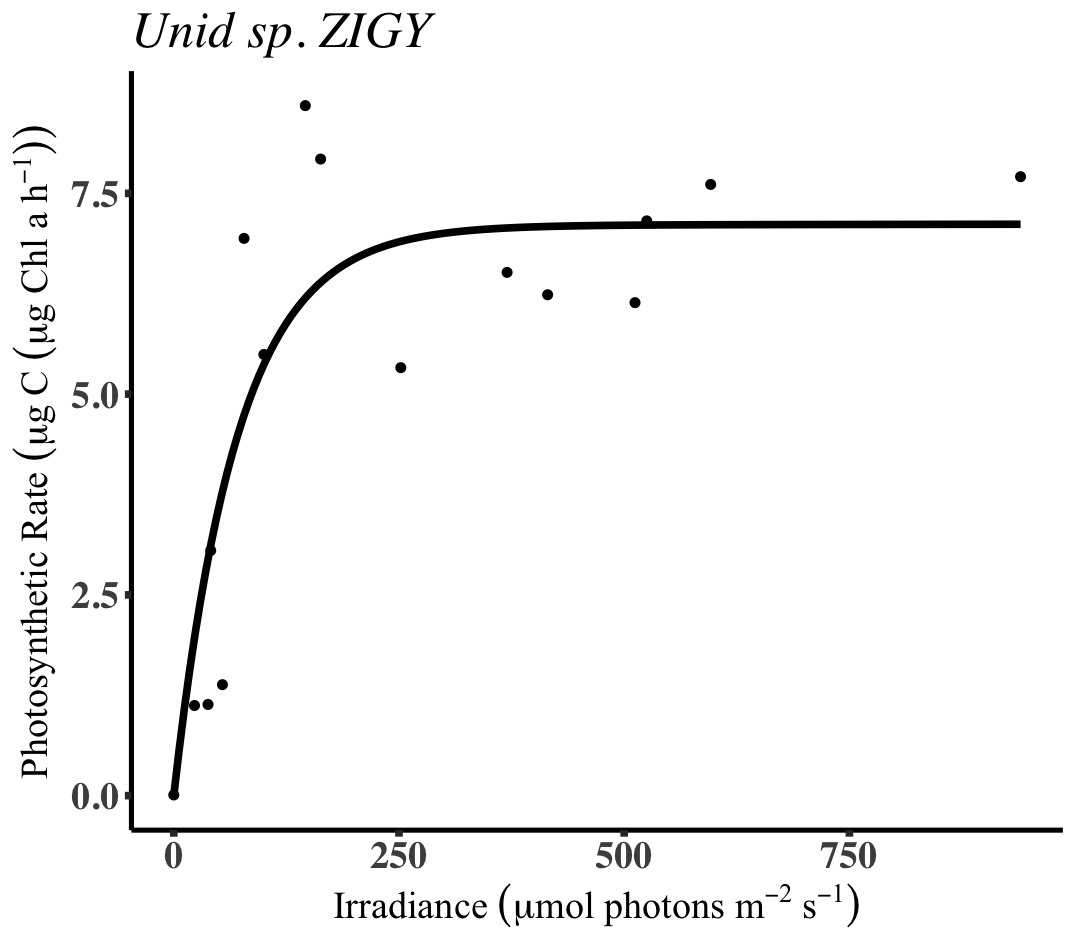

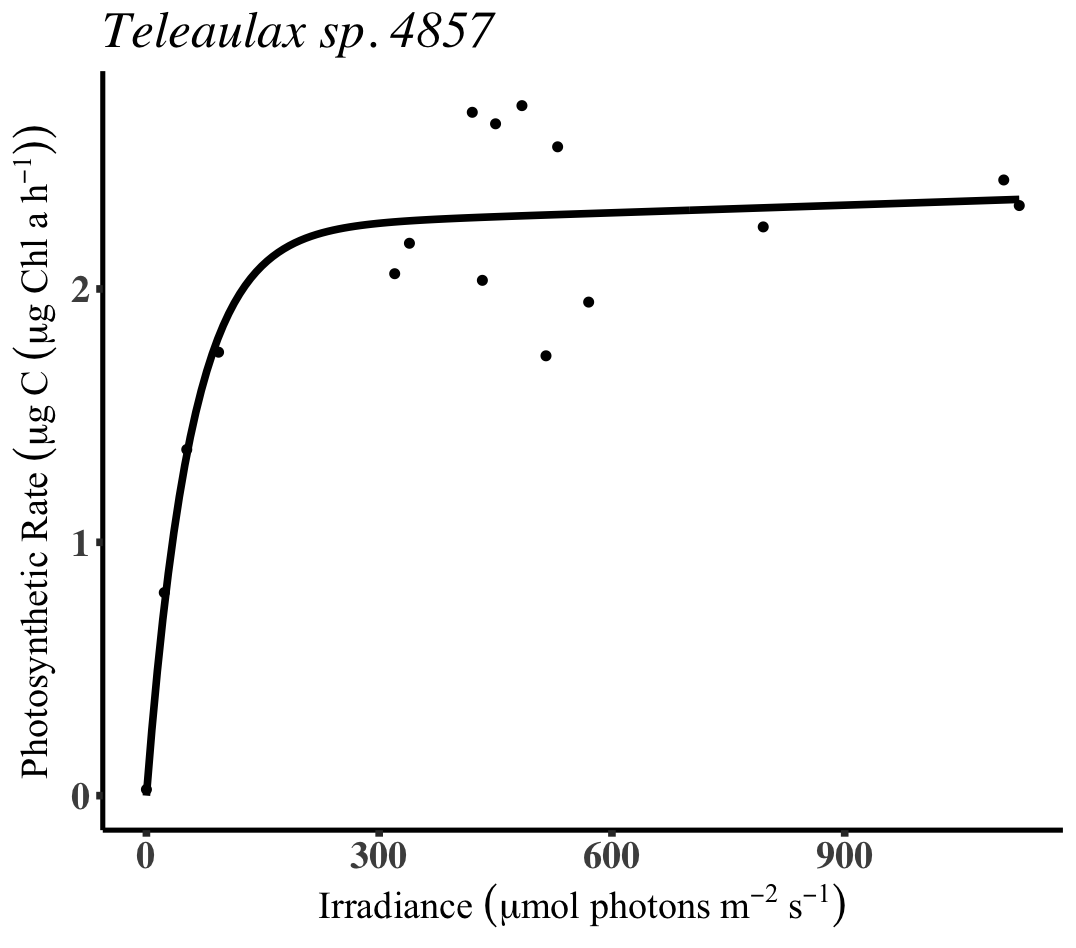

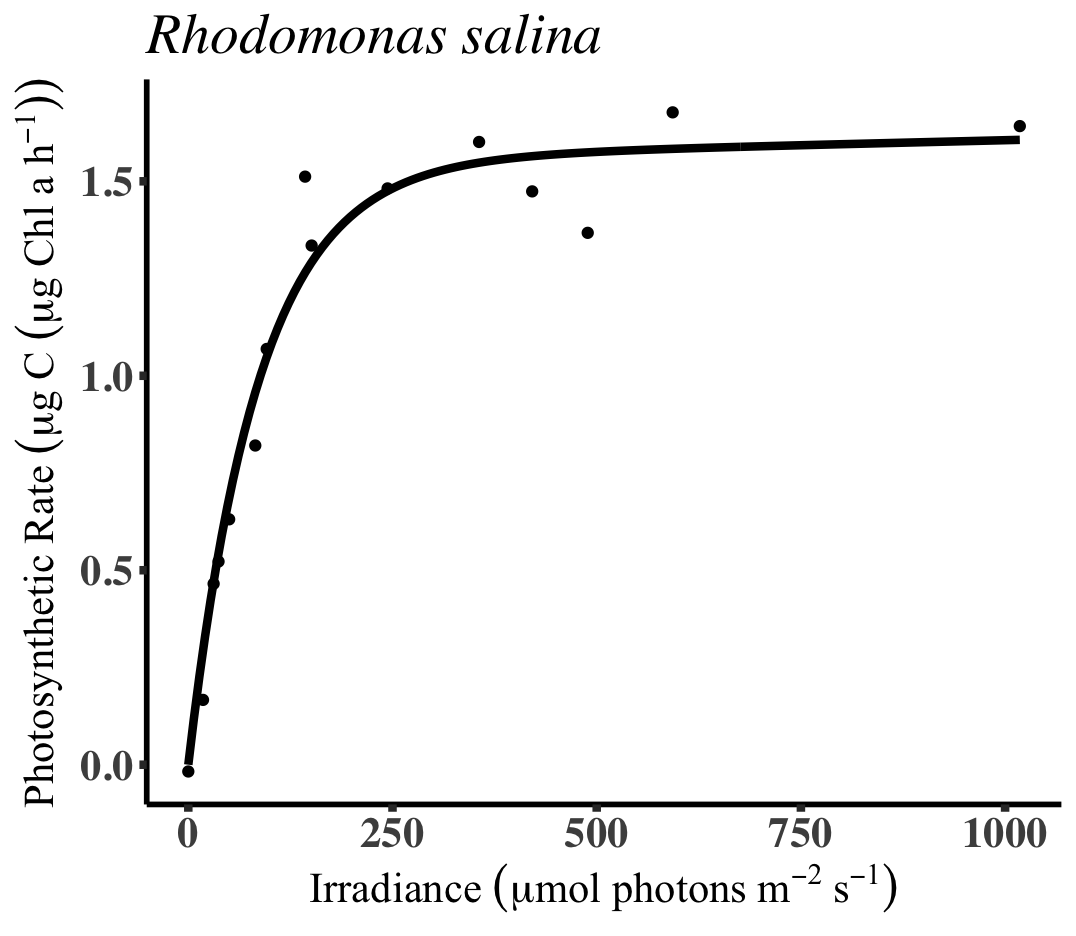

**n)**

**m)**

**o)**
